# Parallel comparison reveals conservation and re-wiring of abiotic stress response networks across three plant species

**DOI:** 10.1101/2024.04.15.589492

**Authors:** Jia Min Lee, Jong Ching Goh, Daniela Mutwil-Anderwald, Marek Mutwil

**Affiliations:** School of Biological Sciences, Nanyang Technological University, 60 Nanyang Drive, Singapore, 637551, Singapore

**Author notes:** Corresponding author: Marek Mutwil, School of Biological Sciences, Nanyang Technological University,60 Nanyang Drive, 637551, Singapore, Singapore.

## Abstract

Abiotic stresses trigger significant alterations in plant morphology, yield, and nutrient composition. To counter these stresses, plants have developed strategies involving gene expression adjustments, impacting their growth and development to ensure survival and reproduction. Although transcriptomic studies have extensively examined stress responses, our understanding of their conservation remains limited. We conducted a comprehensive study on the effects of 24 different environmental and nutrient conditions on the growth yield of three hydroponically grown leafy crops (cai xin, lettuce, and spinach). This research identified optimal conditions for maximizing yield and vitamin content and pinpointed conditions that induce abiotic stress. Our findings highlight the substantial impact of stress on various plant functions, with photosynthesis playing a pivotal role in vitamin content regulation. A detailed comparative analysis of gene expression across these crops revealed conserved gene sets potentially crucial for abiotic stress response. Furthermore, we introduced a novel method that merges regression and genomics to construct reliable gene regulatory networks for stress response. Despite significant conservation of these networks among the studied crops, their key transcription factors showed limited similarity to those in *Arabidopsis thaliana*, suggesting notable functional divergence. To give easy access to the data and comparative transcriptomic tools, we established StressPlantTools, an public online database (https://stress.sbs.ntu.edu.sg/), facilitating further research into stress-responsive genes.

## Introduction

With changing dynamics in global food markets and an expanding population, more research is needed for more resilient food production in urban environments. Hydroponics have emerged as a potential solution for urban farming (Martin and Molin 2019), as this technology can be deployed on rooftops and indoors, and allows controlled light, temperature and nutrients levels for maintaining high growth rates, with the additional advantage of saving water. Thus, in the light of food security, independence of soil quality and local climate, hydroponics techniques have become a major part of global agriculture, in particular for leafy greens (Rajaseger et al. 2023). In combination with ‘smart farming’ which uses sensors and other control systems to constantly monitor nutrient levels and plant vitality, hydroponics can produce up to 20 times the yield per acre of soil-planted crops, e.g. for lettuce, with only 1/20th the amount of water (Majid et al. 2021; Gul and Bora 2023). However, many small to medium-sized farmers commonly use a standard growth medium for practical reasons or because the individual optimal growth conditions of different species are unknown. Consequently, hydroponically grown crops can give low yield and/or decreased nutritional values due to abiotic stress. Standard growth media that are not tailored to the specific plants’ needs may also result in excess waste generated, containing high amounts of unabsorbed nitrate or phosphate, which can pollute the environment including the groundwater (Sutton et al. 2011).

Abiotic stresses can affect hydroponically grown plants, e.g. if nutrient levels are too high or too low, or if temperature and light conditions are not suited to the plants’ growth rates. High temperatures greatly decrease efficiency of photosynthesis and respiration due to changes in membrane fluidity and permeability (Ding et al. 2020; Zhao et al. 2020). Heat also leads to increase in ROS generation from the photosystems which in turn induces lipid peroxidation, inactivating enzymes and degrading proteins (Zhao et al. 2020)). Low levels of macronutrients, such as nitrogen (N), phosphorus (P) and potassium (K), lead to typical deficiency symptoms in plants (van Maarschalkerweerd and Husted 2015). As an essential component of DNA, RNA, proteins and chlorophyll, N deficiency leads to drastic changes in morphology and metabolism, reflected in stunted growth, small leaves and chlorosis (de Bang et al. 2021). NO3-also acts as a signaling molecule, and plays a central role in protecting plants from various environmental stresses (Wang et al. 2018; Khan et al. 2023). P is essential in nucleic acids (DNA, RNA) and phospholipids in cell membranes and central for protein phosphorylation. Low P has a profound impact on energy metabolism (ATP, NADPH), and thus photosynthesis and respiration (de Bang et al. 2021). Physiological functions of K include stomatal regulation, photosynthesis and water uptake (Johnson et al. 2022). K deficiency activates a range of sensing and signaling systems in plants, which involve ROS, Ca2+, phytohormones and microRNAs (de Bang et al. 2021). Increasing light intensities in a physiological range up to 300 uE typically result in better growth and higher soluble sugar and protein content in leafy vegetables. However, high light can affect leaf morphology and lead to leaf curling and/or tipburn (Miao et al. 2023). At the same time, abiotic stresses can cause a change of health-promoting vitamins, as exemplified by vitamin B deficiency under stress (Hanson et al. 2016). Due to the diverse genetic background of the different plant species, it is likely that the sensitivity to abiotic stresses, vitamin levels and the gene regulatory programs controlling the responses to abiotic stresses will differ, but the extent of these differences have not been extensively studied across multiple species.

Plants have evolved mechanisms to perceive abiotic stresses and make appropriate adjustments in their gene expression, and consequently growth and development in order to survive and reproduce (Gong et al. 2020). Transcriptomics has been widely used to study stress (Hirayama and Shinozaki 2010), but insights into the conservation of stress responses are still quite limited. Therefore, we carried out a systematic investigation of 24 different environmental and nutrient conditions on the growth yield of three leafy crop plants that are often grown in hydroponics (cai xin, lettuce and spinach). This allowed us to identify the optimal conditions for each crop for maximum yield and vitamin levels, as well as the conditions leading to abiotic stress. We observed that the stresses have a strong effect on multiple biological pathways, and that photosynthesis is likely a major driver of vitamin levels. We then performed an in-depth parallel analysis of gene expression in the three plants and identified sets of conserved, high-confidence genes that are likely involved in responding to abiotic stress. To build high-confidence gene regulatory networks controlling stress responses, we developed a new approach that combines regression and genomic approaches. Surprisingly, while we observed a significant conservation of the gene regulatory networks across the three crop species, the comparison of the key transcription factors to their *Arabidopsis thaliana* counterparts indicates poor conservation, indicating substantial functional differences between transcription factors across species. Finally, we also established an online stress database of gene expression profiles for the three crops (https://stress.sbs.ntu.edu.sg/) that will allow researchers to carry out comparative analyses driving the discovery of stress responsive genes.

## Methods

### Growth conditions and chambers

We used Aspara® Nature+ Smart Growers (Growgreen Ltd. Hong Kong) which were placed in a MT-313 Plant Growth Chamber (HiPoint, Taiwan), a PGC-9 series controlled environment chamber (Percival Scientific, Inc, Perry, US) or at the ambient conditions of 23-24°C (for cai xin and lettuce), and 22°C (for spinach) in different labs (Figure S1).

### Germination of cai xin and lettuce

Two to three seeds were placed in each seed holder of the Aspara® unit and covered with a germination dome. The tank was filled with tap water, and the unit was switched on to circulate the water. Light conditions during germination were white light for 24 h (40 μmol·m^−2^·s^−1^), and the temperature was 23-24°C. After 3 days, the radicle appeared in most of the seeds (=DAG 0). On DAG 2, one seedling per seed holder was selected, and those with slower growth removed, so in total there were 8 seedlings per Aspara unit. The Aspara® Smart Grower Hydroponic System is based on an ebb-and-flow system and holds 2 L of medium.

### Germination of spinach (Spinacia oleracea var Carmel)

Spinach seeds were sown on cotton balls, kept in the dark, and sprayed regularly with water to keep seeds damp. Within 3 days, 80% of seedlings germinated, i.e., radicles were visible on the seed coat (=DAG 0). They were kept in the dark for an additional 2 days until cotyledons were visible. On DAG 2, 8 seedlings were transferred to the seed holders in the Aspara units. The tank was filled with tap water, and the unit was switched on to circulate the water. Light conditions were white light for 24 h (40 μmol·m^−2^·s^−1^), and the temperature was 23-24°C. One unit was used per growth condition holding 8 plants, and on DAG 12, five plants were selected to ensure sufficient growth space during the last nine days.

### Growth medium

The composition of the half-strength Hoagland’s solution was KH_2_PO4 (0.5 mM), KNO_3_ (3 mM), Ca(NO_3_)_2_ x 4 H_2_O (2 mM), and MgSO_4_ x 7 H_2_O (1 mM). Of the micronutrient stock solution (1000x), 0.5 ml was added to 1L of half-strength Hoagland’s Solution. One liter of micronutrients stock solution contained H_3_BO_3_ (2.86 g), MnCl_2_ x 4 H_2_O (1.81 g), ZnSO_4_ x 7 H_2_O (0.22 g), CuSO_4_ x 5 H_2_O (0.08 g), Na_2_MoO_4_ x 2 H_2_O (0.025 g), CoCl_2_ x 6 H_2_O (0.025 g). One liter of the chelated iron stock solution (200x) contained FeSO_4_ x 7H_2_O (5.56 g) and Na_2_EDTA (7.45 g). 2.5 mL was added to the half-strength Hoagland’s solution. KOH was used to adjust the pH to 5.5. Growth medium levels were checked daily and were replenished regularly in each unit to ensure a stable water level. Furthermore, pH and EC (1.3 dSm^−1^) were monitored every 2-3 days and adjusted by exchanging with fresh growth medium. Plants were harvested on DAG 21.

### Growth conditions during stress experiments

On DAG 5 stress and control conditions were started in half-strength Hoagland’s solution. For nutrient stress conditions, the growth medium was modified (see below). For cai xin and lettuce, control conditions were a light intensity of 202.5 μmol·m^−2^·s^−1^, an R:B:W ratio of 4:1:1, and a light period of 20h at 25°C. For spinach, the control conditions were a light intensity of 130 μmol·m^−2^·s^−1^, an R:B:W ratio of 4:1:1 and a light period of 15h at 22°C.

For the light intensity experiment, cai xin and lettuce were grown at 67, 135, 202.5, and 268 μmol·m^−2^·s^−1^ at 16h light and 25°C. Spinach was grown at 65, 130, 200, and 260 μmol·m^−2^·s^−1^at 15h light and 22°C. For the day length experiment, cai xin and lettuce were grown at 8h, 12h, 20h, and 24h light at 25°C, 200 μmol·m^−2^·s^−1^ and R:B:W 4:1:1. For spinach, the day lengths were 8h, 13h, 18h, 24h light at 22°C, 130 μmol·m^−2^·s^−1^ and R:B:W 4:1:1. For the light quality experiments, the R:B:W ratios were 4:1:1, 4:1:0, 3:1:1 and 3:1:0 for all plant species under otherwise control conditions. For the temperature experiments, the temperatures were 20°C, 25°C, 30°C and 35°C for all plant species under otherwise control conditions.

In the modified N solution, KNO_3_ and Ca(NO_3_)_2_ were replaced with KCl and CaCI_2_, respectively, to give the same concentrations of K and Ca as in the original solutions. In the modified P solution, KH_2_PO4 was replaced by KCl, and in the modified K solution, KH_2_PO4 and KNO_3_ were replaced by NaH_2_PO4 and NaNO_3_.

Modified N Solution
|  | 0% N (mM) | 25% N (mM) | 50% N (mM) | 100% N (mM) | 150% N (mM) |
| --- | --- | --- | --- | --- | --- |
| $\text{KNO}_3$ | 0 | 0.75 | 1.5 | 3 | 4.5 |
| KCl | 3 | 2.25 | 1.5 | 0 | 0 |
| $\text{Ca}(\text{NO}_3)_2$ | 0 | 0.5 | 1 | 2 | 3 |
| $\text{CaCl}_2$ | 2 | 1.5 | 1 | 0 | 0 |

Modified P solution
|  | 0% P (mM) | 25% P (mM) | 50% P (mM) | 100% P (mM) | 150% P (mM) |
| --- | --- | --- | --- | --- | --- |
| $\text{KH}_2\text{PO}_4$ | 0 | 0.125 | 0.25 | 0.5 | 0.75 |
| KCl | 0.5 | 0.375 | 0.25 | 0 | 0 |

Modified K Solution
|  | 0% K (mM) | 25% K (mM) | 50% K (mM) | 100% K (mM) | 150% K (mM) |
| --- | --- | --- | --- | --- | --- |
| KH<sub>2</sub>PO<sub>4</sub> | 0 | 0.125 | 0.25 | 0.5 | 0.75 |
| NaH<sub>2</sub>PO<sub>4</sub> | 0.5 | 0.375 | 0.25 | 0 | 0 |
| KNO<sub>3</sub> | 0 | 0.75 | 1.5 | 3 | 4.5 |
| NaNO<sub>3</sub> | 3 | 2.25 | 1.5 | 0 | 0 |

NaOH was used to adjust the pH to 5.5.

### Sampling of plants and determination of fresh weight

On DAG 12, five of eight seedlings were selected based on healthy conditions and similar growth. On DAG 21, out of five biological replicates per condition, the three most healthy biological replicates similar in growth were selected. Of these, three to four mature leaves were cut, weighed (minimum total weight 0.5 g) and immediately flash-frozen in liquid nitrogen, and stored at −80°C. After sampling for RNA, the rest of the leaves were cut, and the fresh weight (FW) was measured to obtain the total fresh weight of each biological replicate.

Student’s two-sided t-tests (Student 1908) were carried out to determine the effects of stress conditions on the growth of the species with respect to the largest mean fresh weight measured across all stress conditions under the respective stress experiment. A stress condition was determined to significantly alter the growth of the species when the p-value (corrected with the Benjamini-Hochberg procedure (Benjamini and Hochberg 1995)) is less than 0.05.

### RNA isolation and sequencing

All leaf samples were ground by mortar and pestle, and the frozen powder was aliquoted and stored at −80°C until use. RNA was isolated from stressed and control plants (three biological replicates each) using Spectrum^TM^ Plant Total RNA Kit (Sigma). Quality control of all the extracted RNA (triplicates for each condition) was done by Novogene (Singapore) using Nanodrop and agarose gel electrophoresis (for purity and integrity) before sample quantitation and further analyses of integrity (Agilent 2100 Bioanalyzer). The library type was a eukaryotic directional mRNA library. Library construction from total RNA, including eukaryotic mRNA enrichment by oligo(dT) beads, library size selection, and PCR enrichment, was performed by Novogene using NEBNext® Ultra™ II Directional RNA Library Prep Kit for Illumina®. The libraries were sequenced with NovaSeq-6000, paired-end sequencing at 150 base pairs and at sequencing depths of approximately 20 million reads per sample (6 Gb/sample).

### Vitamin measurements

LCMS grade ethanol (90%) was added to 100 mg of frozen leaf powder to give a concentration of 100 mg/ml. The samples were vortexed for a few minutes before sonicating in ice water for 10 minutes, followed by centrifuging at 14,000 rpm for 10 minutes. The supernatant was collected and injected into LCMS for analysis. Quality control samples were prepared by pooling the extracts, which were analyzed along with the samples to ensure instrument stability.

Quantification of water-soluble vitamins was done using a Waters HSS T3 column (2.1mm x 100mm, 1.8µm) with Mobile Phase A (Water + 0.1% formic acid) and Mobile Phase B (Acetonitrile + 0.1% formic acid) at a Column Temperature of 40°C and a Sample Temperature of 4°C. MS analysis was done using a Xevo TQ-S machine (Waters), Ion mode: ESI positive, Acquisition mode: MRM, Capillary Voltage: 1.5 kV, Desolvation temperature: 600°C, Cone gas flow: 150 L/h, Desolvation gas flow: 1200 L/h, and Source temperature: 150°C.

Standard solutions were prepared in LCMS grade methanol to provide a series of standard solutions with gradient concentrations (10-10000 ng/ml) to make the calibration curves.

### Gene expression and differential gene expression estimation

Quantification of the abundances of transcripts from RNA sequencing data was carried out with kallisto v0.46.1 (Bray et al. 2016). The RNA sequencing data of cai xin, lettuce, and spinach were pseudoaligned against *Brassica rapa* (BRADv1.2 (Cheng et al. 2011; Chen et al. 2022), Phytozome (Goodstein et al. 2012)), *Lactuca sativa* (V8 (Reyes-Chin-Wo et al. 2017), Phytozome), and *Spinacia oleracea* (Spov3 (Hulse-Kemp et al. 2021), Phytozome) CDSs, respectively. Gene expression output from kallisto includes count and TPM (transcripts per million). Based on the TPM, the Pearson correlation coefficient (PCC) was computed between all replicates and stress conditions. Clustermap was adopted for each species to illustrate the correlation, and similarities were analyzed according to the Euclidean distances between the samples.

Differential gene expression was determined using DESeq2 v1.40.1 (Love et al. 2014) with the count outputs from kallisto. For the various stress conditions, they were compared with their respective stress experiment control. Genes were identified as differentially expressed at |*log*_2_*FC*| ≥ 1 and adjusted p-value ≤ 0.05 (corrected with the Benjamini-Hochberg procedure).

### Gene annotation and differentially expressed biological functions

The biological function annotations of the genes of all three species were assigned with Mercator4 v5.0 (Schwacke et al. 2019), where an annotation refers to a MapMan bin. Transcription factors (TFs) and their corresponding gene symbols for cai xin and lettuce were identified with iTAK v1.6 (Zheng et al. 2016), while for spinach, they were identified with iTAK v1.7.

Survival function was used to identify the significantly differentially expressed biological functions, where significance was with the condition that the resultant p-value corrected with the Benjamini-Hochberg procedure was less than 0.05. For inference of similarities between the biological functions (rows) and stress conditions (columns), Jaccard distances (JDs) were computed between the biological functions and stress conditions, respectively. The JDs were used to carry out the hierarchical clustering of the biological functions and stress conditions.

### Identification of vitamin biosynthetic genes and pathways and evaluation of the effects of stress conditions on vitamin levels

Plant Metabolic Network (PMN) database (Hawkins et al. 2021) was used to identify the vitamin biosynthetic pathways and their corresponding Enzyme Commision (EC) numbers. The biosynthetic pathways of vitamin B3 (nicotinamide and nicotinic acid), vitamin B5 (pantothenic acid), vitamin B6 (pyridoxal and pyridoxine), riboflavin and thiamine were identified as nicotine biosynthesis, phosphopantothenate biosynthesis I, pyridoxal 5’-phosphate biosynthesis II, flavin biosynthesis I (bacteria and plants), and thiamine diphosphate biosynthesis IV (eukaryotes), respectively. Ensemble Enzyme Prediction Pipeline (E2P2 v4.0) (Chae et al. 2014) was employed on the protein sequences of the three species for functional annotations with EC numbers (not exclusively).

The genes involved in the synthesis of the various vitamins were identified through the EC numbers associated with the vitamin biosynthetic pathways.The Spearman rank-order and Pearson correlation coefficients were computed with the gene expression (TPM) and vitamin level datasets. Survival function was applied to identify the biological functions significantly associated with each vitamin, based on the top 50 positively and negatively correlated genes corresponding to each vitamin. This approach aimed to highlight potential relationships between biological functions and vitamin levels.

Student’s two-sided t-tests were carried out on the vitamin levels across various stress conditions with respect to the vitamin level of control conditions. A stress condition was determined to significantly regulate a vitamin when the p-value (corrected with the Benjamini-Hochberg procedure) is less than 0.05, and the corresponding fold change was computed as indication of the magnitude of change.

### Detection of orthogroups and calculation of significant similarities between stresses

Homologous relations between cai xin, lettuce and spinach were inferred with OrthoFinder v2.5.5 (Emms and Kelly 2015, 2019), through which differentially expressed orthogroups can be identified by mapping the differentially expressed genes (DEGs) to their orthogroups. Similarities between the responses of all three species to different stress conditions were measured with the Jaccard index (JI). For stress conditions comparison of a species, JI was computed between all stress conditions with the DEGs datasets. For across-species comparison, JI was computed between all stress conditions and species based on the differentially expressed orthogroups. The significance of the quantified similarities was evaluated through permutation analysis, where the observed JIs were compared with the permuted JIs. A thousand permutations were executed for an observed JI, and significance was assumed when the resultant p-value (corrected with the Benjamini-Hochberg procedure) was less than 0.05.

A conserved orthogroup was defined as an orthogroup identified in all species that were differentially expressed in a similar manner (up- or down-regulated) under the same surplus or deficient stress condition, where surplus was an increase in stress parameter with respect to the control condition and deficient was the inverse.

### Construction of gene regulatory networks

GENIE3 (Huynh-Thu et al. 2010; Aibar et al. 2017) was employed to construct the gene regulatory networks (GRNs) for individual species based on the TPM expression of DEGs and the regulators were specified as the TFs. For high confidence networks, the leading 5% (by the ranks of non-zero weight) TF-target edges were used for downstream analyses (Figure 5D). The relationships between the TFs and their targets were assessed through the PCC analysis. Positive correlations were indicative of TFs acting as activators, while negative correlations suggested TFs acting as repressors.

Stress condition-specific GRNs were constructed with conserved TF-target edges. Similar to the identification of conserved orthogroups, TF-target edges were first mapped to their respective orthogroups for across-species comparison. Conserved TF-target edges were defined as edges differentially expressed in a similar manner (up- or down-regulated) under the same surplus or deficient stress condition in > 1 species (Figure 5D). The significance of the conserved edges was evaluated through permutation analysis, where the observed conserved edges were compared to the permuted conserved edges. Enrichment score was computed based on the ratio between the observed conserved edges and the mean of the permuted conserved edges. Significance was assumed when p-value (corrected with the Benjamini-Hochberg procedure) was < 0.05 and enrichment score > 1. Henceforth, we define TFs of the significantly conserved TF-target edges as ‘the stress condition-specific conserved TFs’.

### Biological functions of stress condition-specific conserved TFs

To gain insight into the biological functions the stress condition-specific conserved TFs were to regulate, the associated biological functions (MapMan bins) of the TF’s targets were determined. The distribution of the targets’ biological functions were normalized by the corresponding MapMan bin sizes. Hierarchical clustering was imposed by calculating the Euclidean distance between the TFs for emphasizing similar TFs.

### Stress condition-specific conserved TF response with experimentally verified gene Arabidopsis genes

The stress condition-specific conserved TFs were queried against *Arabidopsis thaliana* (Araport11 (Cheng et al. 2017), TAIR (The Arabidopsis Information Resource (TAIR))) with BLAST (Camacho et al. 2009), a local similarity bioinformatics tool, for their corresponding best hits. The stress conditions the conserved TFs (columns) were differentially expressed under were mapped against the experimentally verified functions of the best BLAST hits (rows), where a match between the stress condition and experimentally verified function would be highlighted in red or blue, and mismatch would be in gray. The columns were grouped according to their species and the stress conditions (surplus and/or deficient) the conserved TFs were found to be differentially expressed. Red and blue (matches) were used as further indications of up- and down-regulation, respectively, responses of the conserved TFs under the corresponding stress condition. The *Arabidopsis thaliana* best BLAST hits (rows) were grouped according to their functions, where the miscellaneous category consists of gene ontology (GO) terms like “response to hydrogen peroxide”, “response to wounding”, “cellular response to hypoxia”, “response to water deprivation”, and other functions that were not represented as a separate category. Repeated occurrences of the same best BLAST hit under different function categories indicate multiple functions.

Permutation analysis was performed to evaluate the significance of the conservation of function between the stress condition-specific conserved TFs predicted role and their *Arabidopsis thaliana* best BLAST hit functions. The observed matches were compared to the permuted matches through a thousand permutations of the stress conditions the conserved TFs were differentially expressed, while keeping the best hits and their experimentally verified functions constant. Significance was assumed when p-value (corrected with the Benjamini-Hochberg procedure) was < 0.05.

### Establishment of StressPlantTools database

Using the CoNekT framework admin panel (Proost and Mutwil 2018), we constructed the database with the generated gene expression data. We employed the Highest Reciprocal Rank metric to build the coexpression networks (Mutwil et al. 2010). For each species, coexpression clusters were obtained by using the Heuristic Cluster Chiseling Algorithm (HCCA). The database is hosted on an Apache server operating under Windows OS.

### Data availability

The raw sequencing data is available at ENA under the accession number: E-MTAB-14018.

## Results

### Abiotic stress experiments for lettuce, cai xin and spinach

To determine the effects of environmental conditions on the growth of cai xin (Figure 1A), lettuce (Figure 1B) and spinach (Figure 1C), plants were grown for 16 days in the respective experimental condition, and the phenotypes and FWs were captured on DAG 21 (Figure 1D). The control medium for all plants was half-strength Hoagland solution, a widely used standard growth medium characterized by high levels of N and K, thus suitable for plants with high nutrient demands. The significance of the observed growth differences was evaluated by Student’s two-sided t-tests with respect to the maximum mean FW (indicated as red bars).

**Figure 1.**
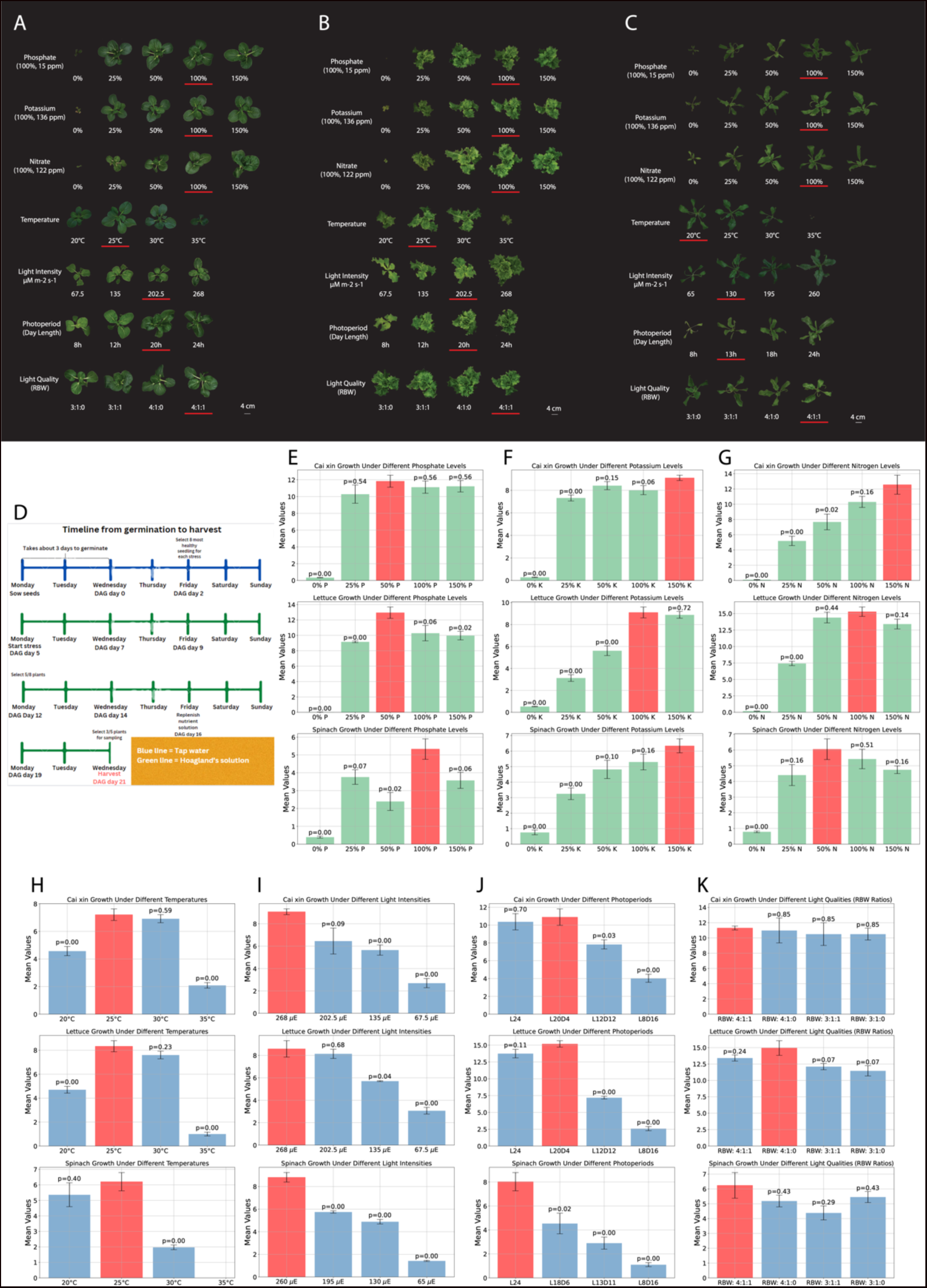
Abiotic stress experiments for hydroponically-grown plants. Phenotype of A) cai xin, B) lettuce, and C) spinach on DAG 21 for phosphate, potassium, nitrate, temperature, light intensity, day length, and light quality stress experiments. Control condition of each stress is highlighted with red underline. D) Overview of the experiment timeline for the experiments. Fresh weights of all three species under all stress conditions were measured on DAG 21. The mean fresh weight data of E) phosphate, F) potassium, G) nitrate, H) temperature, I) light intensity, J) photoperiod, and K) light quality stress experiments are presented as bar charts. Error bars are used to indicate the standard deviation. Student’s two-sided t-test p-values (corrected with the Benjamini-Hochberg procedure) are also annotated for each stress condition with respect to the maximum mean fresh weight (red bars).

Complete removal of a macronutrient from the growth medium, either 0% N, 0% P, or 0% K, resulted in severely reduced growth of all three species (Figures 1E-G). In media with reduced macronutrients (25%, 50%) for P and K (Figures 1E-F), cai xin displayed similar FW amounts as compared to control (100%). In contrast, it showed a concentration-dependent increase in FW at increasing N levels up to 100%, yielding a maximum of 12g (Figure 1G). This indicates that cai xin is dependent on high N (but not P or K) for better growth. Lettuce grew well in lower P levels (25%, 50%), but showed decreased growth under low K (25%, 50%) and low N (25%). Spinach grew better with increasing K concentrations (Figure 1F), but shows varying FW at different P and N levels. Increasing the concentration of the nutrients (N, P and K) to 150% did not result in a significantly enhanced growth phenotype for any of the plants. Overall, this shows that the three crops have different requirements in macronutrients for optimal FW.

Temperature plays an important role for obtaining maximum FW. Both cai xin and lettuce grow well at 25°C and 30°C, but show reduced growth at 20°C and 35°C (Figure 1H). Spinach grows best at 25°C, but is more sensitive to heat at 30°C and 35°C. While cai xin and lettuce could still grow at 35°C, spinach plants died after a few days (Figure 1H).

Low light intensities (65 to 135 μmol·m^−2^·s^−1^) and shorter photoperiod (8 to 13 hours of light) significantly decreased the growth of all three species compared to the controls. In cai xin and lettuce, the highest light intensity (268 μmol·m^−2^·s^−1^) and longest photoperiod (24 h light) did not promote further growth (Figures 1H-I). In contrast, spinach showed maximum FW at the highest light intensity (260 μmol·m^−2^·s^−1^) or the longest photoperiod (24 h light). However, some leaves were yellow at the tips and displayed curling (data not shown). Thus, a lower light intensity of 130 μmol·m^−2^·s^−1^ and a shorter photoperiod of 13 h light was chosen for spinach as default for the other experiments, which is in line with previous reports (Zou et al. 2020). Variation in light quality by modifying rations of red, blue and white light did not promote significant changes to the phenotype in all three species (Figures 1J-K).

### Gene expression analysis of abiotic stress responses, and establishment of stress database

To better understand how the three species respond to the different growth conditions on the level of gene expression, we performed RNA-sequencing. For each of the species and stress conditions, gene expression data were generated in triplicates (Supplemental Data 1-3 for expression matrices of cai xin, lettuce and spinach, respectively). A total of 276 RNA-seq samples for 31 stress conditions (30 for spinach due to death at 35°C) were obtained from all three species. Clustering of the samples revealed a high similarity across the replicates, and unique clustering of deficiencies in N, P and K of cai xin (Figure S2), lettuce (Figure S3) and spinach (Figure S4), indicating a strong transcriptional response to these stresses.

To determine the genes that exhibit differential expression in various stress conditions, DEGs were identified with DESeq2 for all stress conditions and species (Figures 2A-C and Tables S3-5). Similar to the growth phenotype observed in Figure 1, across the three species, deficiencies in N, P, K, photoperiod and high temperature resulted in significantly large changes to the gene expressions, while light quality had little effect on the gene expressions. Lettuce exhibited higher sensitivity to the low light (Figure 2B). Across species, stress condition 0% K demonstrated consistently a large number of DEGs.

**Figure 2.**
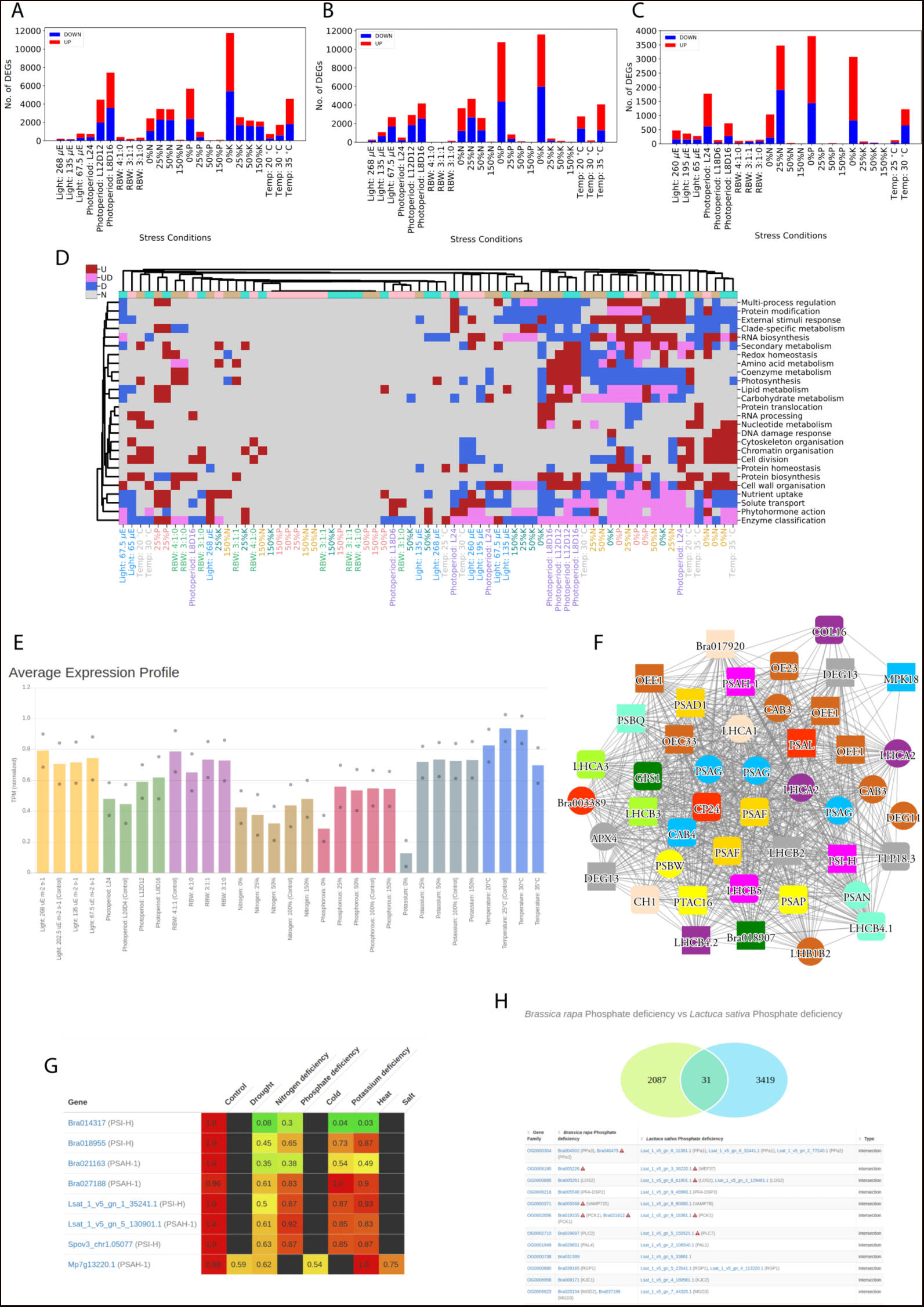
Expression profile and co-expression analyses of stress responses. Differential gene expression analysis of A) cai xin, B) lettuce and C) spinach. The red and blue bars represent up- and down-regulated genes, respectively. The x-axis represents different stress conditions while the y-axis shows the number of differentially expressed genes (DEGs) for a given stress. D) Significantly (p-adj < 0.05) up- (red), down-regulated (blue) or both (pink) biological functions for the different stress conditions and species. The different stress conditions are in columns, while the biological functions are in rows. Only biological functions that were differentially expressed in three or more columns are shown. Similarities between the responses were highlighted by clustering the columns and biological functions across all three species, and the columns are color-coded such that tan, turquoise and pink represent cai xin, lettuce and spinach, respectively. The labels of the stress condition are also colored according to their stress experiments. E) Average expression profiles in the photosynthetic cluster. The colors indicate different stress experiments (x-axis), while the average expression is shown on the y-axis. F) Co-expression network of genes found in the photosynthetic cluster. Nodes represent genes, edges connect co-expressed genes, while colored shapes represent different orthogroups. G) Expression values of PSI-H genes in cai xin, lettuce, spinach and *Marchantia polymorpha*. The genes are shown in rows, while the stresses and controls in columns. H) Venn diagram containing the genes that show conserved upregulation in phosphate deficiency in cai xin and lettuce.

To gain insight into the biological functions that were affected by the stress conditions, significantly differentially expressed biological functions were determined from the DEGs. Results revealed high similarities between the N, P and K deficiencies across all species, with coenzyme metabolism and photosynthesis being significantly downregulated, solute transport, enzyme classification and phytohormone action being both significantly up- and down-regulated, and protein modification and RNA biosynthesis being significantly upregulated, indicating conserved responses (Figure 2D). In particular, under 0% N stress condition, cytoskeleton organisation, cell division, chromatin organisation, cell wall organisation were upregulated and external stimuli response was downregulated for all species. A level 2 analysis, conducted with the second level of (sub-) MapMan bins (Figure S5), was in agreement with previous analysis, whereby N, P and K deficiencies showed high similarity across all species. At 25% N stress condition, transcriptional regulation was significantly upregulated, carrier-mediated transport and EC_1 oxidoreductases were both significantly up- and down-regulated, and photophosphorylation, calvin cycle, oxidative pentose phosphate pathway, chlorophyll metabolism, DNA replication, pectin and sulfur assimilation were some of the significantly downregulated biological functions for all species.

The gene expression data for all the stress experiments on the three species are made available on a stress.plant.tools database (https://stress.sbs.ntu.edu.sg/), along with the data for *Marchantia polymorpha* (Tan et al. 2023).

The database serves as a platform for visualization of expression profiles, co-expression networks and various comparative analyses. To demonstrate the utility of the database, we analyzed a co-expression cluster containing photosynthetic genes. Average expression profile (Figure 2E) revealed that the expression of the photosynthetic genes were lower under N, P, and K deficiency stress conditions, which is in agreement to that observed in Figure 2D. Meaningful co-expression networks can also be illustrated, where an example of the cluster consisted of many known genes important for photosynthesis (Figure 2F). Comparative heatmap allows the depiction of the expression of multiple genes across stress and species for comparison purposes. An example of the photosystem I subunit H protein showed downregulation across all stresses, especially so during nitrogen deficiency (Figure 2G). Genes demonstrated conserved responses can be determined with the ‘Compare specificities’ tool. For cai xin and lettuce subjected to phosphate deficiency, 31 orthogroups were found to be shown conserved responses (Figure 2H). By cross-referencing the conserved orthogroups’ gene symbols with *Arabidopsis thaliana*, six orthogroups (~20% of the conserved orthogroups) had been experimentally verified to have distinct roles under phosphate deficiency conditions (Table 1). While the function of the other genes in the list is not yet related to phosphate deficiency, their conservation strongly suggests their function in responding to this stress. In conclusion, the stress.plant.tools database allows the exploration of stress-specific expression profiles and will be an invaluable platform for studies of stresses.

**Table 1.**
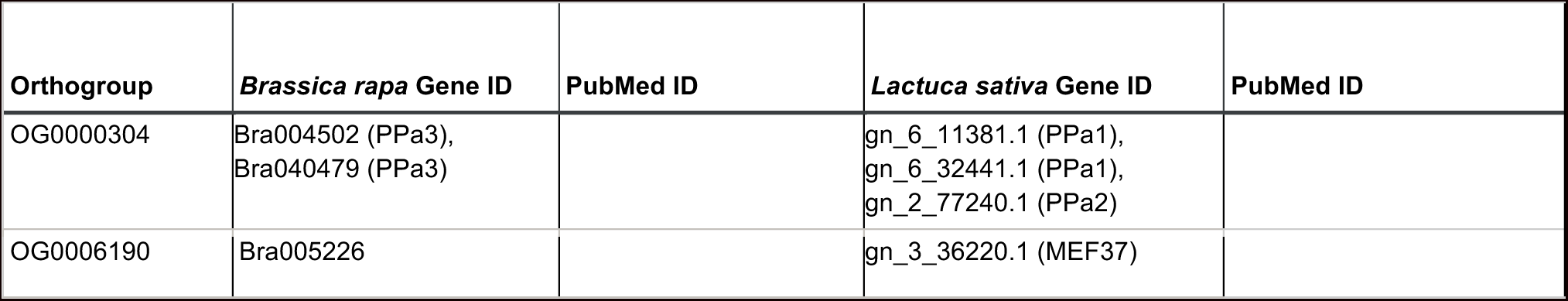
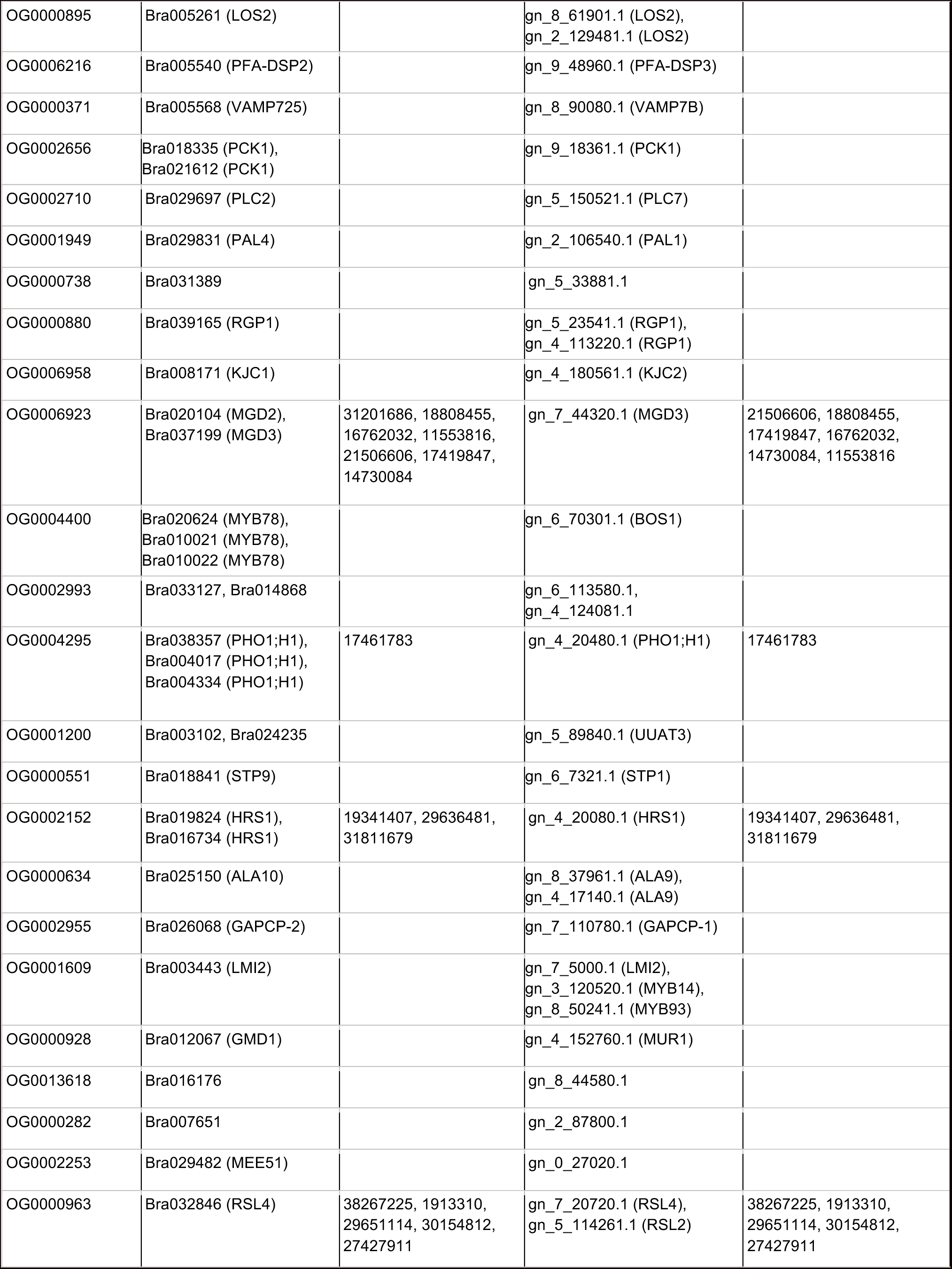
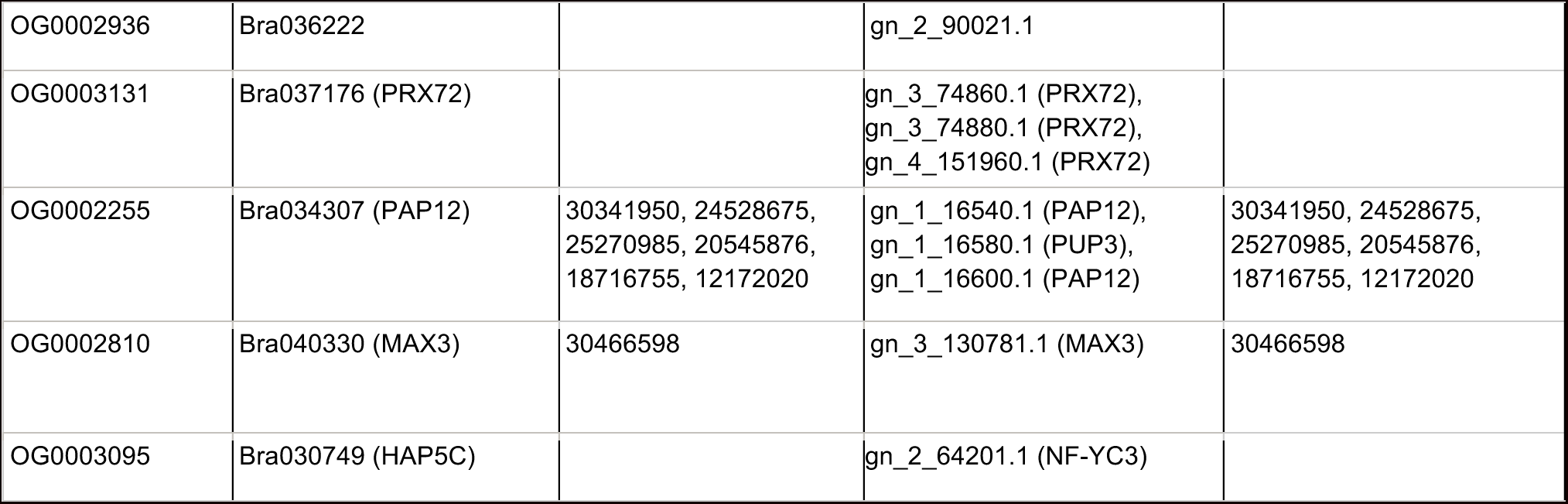
Phosphate starvation-specific orthogroups for *Brassica rapa* and *Lactuca sativa*. For brevity, lettuce gene IDs have been shortened.

### Vitamin levels are not correlated with the expression of biosynthetic genes

To analyze the effects of the various stress conditions on vitamin levels, we measured two forms of vitamin B3, which are nicotinamide (Figure S6) and nicotinic acid (Figure S7), pantothenic acid (vitamin B5) (Figure S8), and two forms of vitamin B6, which are pyridoxal (Figure S9) and pyridoxine (Figure S10), riboflavin (Figure S11) and thiamine (Figure S12) across the stress conditions (Table S6). In general, cai xin contains the highest levels of vitamins among the three species, while spinach contains the least (Table S6). Notably, this disparity is prominent in levels of vitamin B3 (Figures S6-S7), where e.g., cai xin contains > 100 pmol/mg of nicotinamide, while spinach typically contains < 30 pmol/mg (Figure S6). The analysis also revealed that certain stress conditions can alter the vitamin levels, such as lower levels of pyridoxal were measured under potassium and nitrogen deficiencies (0% K and 25% N, respectively) in lettuce (Figure 3A). Examination of the heatmap depicting the normalized TPM of pyridoxal biosynthetic genes in lettuce across different stress conditions (Figure 3A) suggested no discernible trend to indicate regulation of vitamin levels on a transcriptional level.

**Figure 3.**
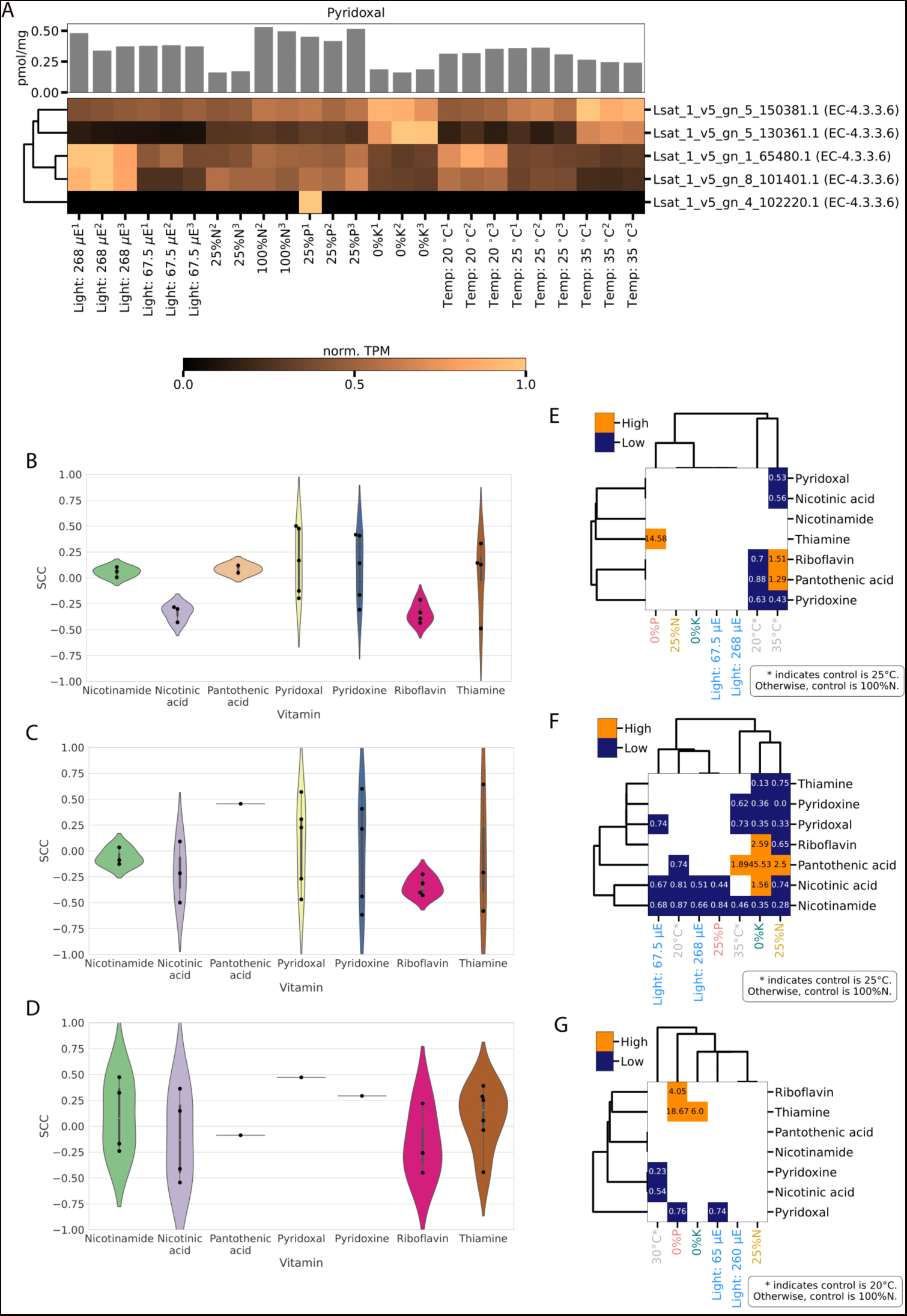
Vitamin levels and expression values of corresponding biosynthetic genes. A) Pyridoxal levels (in pmol/mg) and row-normalized TPM of the last step biosynthetic enzymes for various stress conditions of lettuce are represented by a bar chart and heatmap, respectively. Replicates are differentiated by superscript numbers. Hierarchical clustering based on Euclidean distance between the biosynthetic enzymes were employed to highlight the similarities between the enzymes. Violin plots depict Spearman’s rank correlation coefficient (SCC), capturing the similarity of expression levels of enzymes and the corresponding vitamins for B) cai xin, C) lettuce, and D) spinach. The heatmap indicates vitamin levels that are significantly (p-adj < 0.05) higher (orange), lower (blue) or non-significant (white) in the different stress conditions with respect to a control condition for E) cai xin, F) lettuce, and G) spinach. Temperature stress conditions are compared against the 25°C stress condition for cai xin and lettuce, and 20°C stress condition for spinach. Non-temperature stress conditions are compared against the 100% N stress condition for all species. The fold change values are annotated within each cell of the heatmap.

To further validate whether vitamin levels are regulated at the transcriptional level of their respective biosynthetic enzymes, we measured correlation between the vitamin levels and the enzymes using Spearman’s rank correlation coefficient (SCC) (Figures 3B-D) and Pearson correlation coefficient (PCC) (Figure S13). Overall, we observed poor correlations between the biosynthetic enzymes’ expressions and vitamin levels (median values around 0, Figures 3B-D), suggesting vitamin biosynthesis is not regulated by changing the transcription of the corresponding biosynthetic genes.

While the biosynthetic enzymes were not strongly correlated with their products, we observed strong correlations of vitamin levels to other genes (Table S7). To better understand their functions, we performed a MapMan enrichment analysis (Figures S14-S15, p-adj < 0.05). Based on both SCC and PCC analyses, photosynthesis and coenzyme metabolism are negatively correlated with pantothenic acid, and RNA processing and photosynthesis are negatively and positively correlated to nicotinic acid, respectively, for cai xin. Photosynthesis is negatively correlated to nicotinic acid and pantothenic acid, while positively correlated to pyridoxal, pyridoxine and nicotinamide for lettuce. Across species, cai xin and lettuce demonstrated conserved negative correlation between photosynthesis and pantothenic acid. Overall, this indicates that the regulation of photosynthesis is likely a major determinant of vitamin levels in plants.

Based on results of the Student’s two-sided t-test, not all stress conditions exhibit a significant (p-adj < 0.05) influence on the levels of vitamins across all species. Moreover, the significance of a stress on the vitamin levels of one species is not indicative of a similar effect on another species. Variations of stress conditions were shown to have less significant effect on cai xin and spinach (Figures 3E and 3G), but a more pronounced effect on lettuce (Figure 3F). Pyridoxal levels were significantly reduced to one-third of the measured for 100% N when lettuce was subjected to potassium and nitrogen deficiencies (Figure 3F), which is in agreement with Figure 3A. Noticeably, potassium deficiency for lettuce also significantly enhanced the levels of riboflavin, nicotinic acid, and particularly pantothenic acid, which was increased by a factor of 45.53. High temperature (35°C), and potassium and nitrogen deficiencies (0% K and 25% N) significantly enhanced the synthesis of pantothenic acid in lettuce, while lower temperature (20°C) weakened the synthesis. Nicotinamide in lettuce was reduced under any changes of stress conditions from the controls. Conserved responses were also identified across the species, such as 0% P stress condition significantly increased the level of thiamine in cai xin and spinach, and high temperature reduced the pyridoxine level in all species. Taken together, abiotic stress can significantly affect, and even increase, vitamin levels, suggesting that a more elaborate growth regime can be used to tailor the metabolic composition of plants.

### Conservation of stress responses across species

To better test whether the stress responses are conserved across stresses and species, we investigated whether the three species show significantly similar gene expression changes with a permutation analysis. The permutation analysis revealed that N, P and K deficiency responses were significantly (p-adj < 0.05) similar across species (Figure 4A, brown rectangles, Tables S8-S11). Surprisingly, the N, P and K responses elicit a similar response. For instance, 0% N to 50% N stress conditions of lettuce elicited similar responses as N, P, and K deficiencies stress conditions in cai xin (Figure 4A, lower left corner, blue cells). For upregulated DEGs, same stress conditions upregulate similar sets of genes, particularly between lettuce and cai xin (upper right triangle, Figure 4A). Similar to the downregulated DEGs, N, P, and K deficiencies also upregulate similar sets of genes across different N, P, and K deficiencies stress conditions. For instance, 0% K stress condition of lettuce upregulated similar sets of genes as 0% N and 0% P stress conditions of cai xin. For spinach, similarities were observed across species, albeit fewer. For within species comparison, conservations were observed between the various stress conditions of each stress experiment (Figure 4A, near main diagonal, yellow and green cells). Notably, strong conservation responses across N, P, and K deficiencies stress conditions were observed for both up- and down-regulated genes. Overall, we conclude that many of the stress responses are conserved, and that N, P, and K deficiencies elicit a similar response.

**Figure 4.**
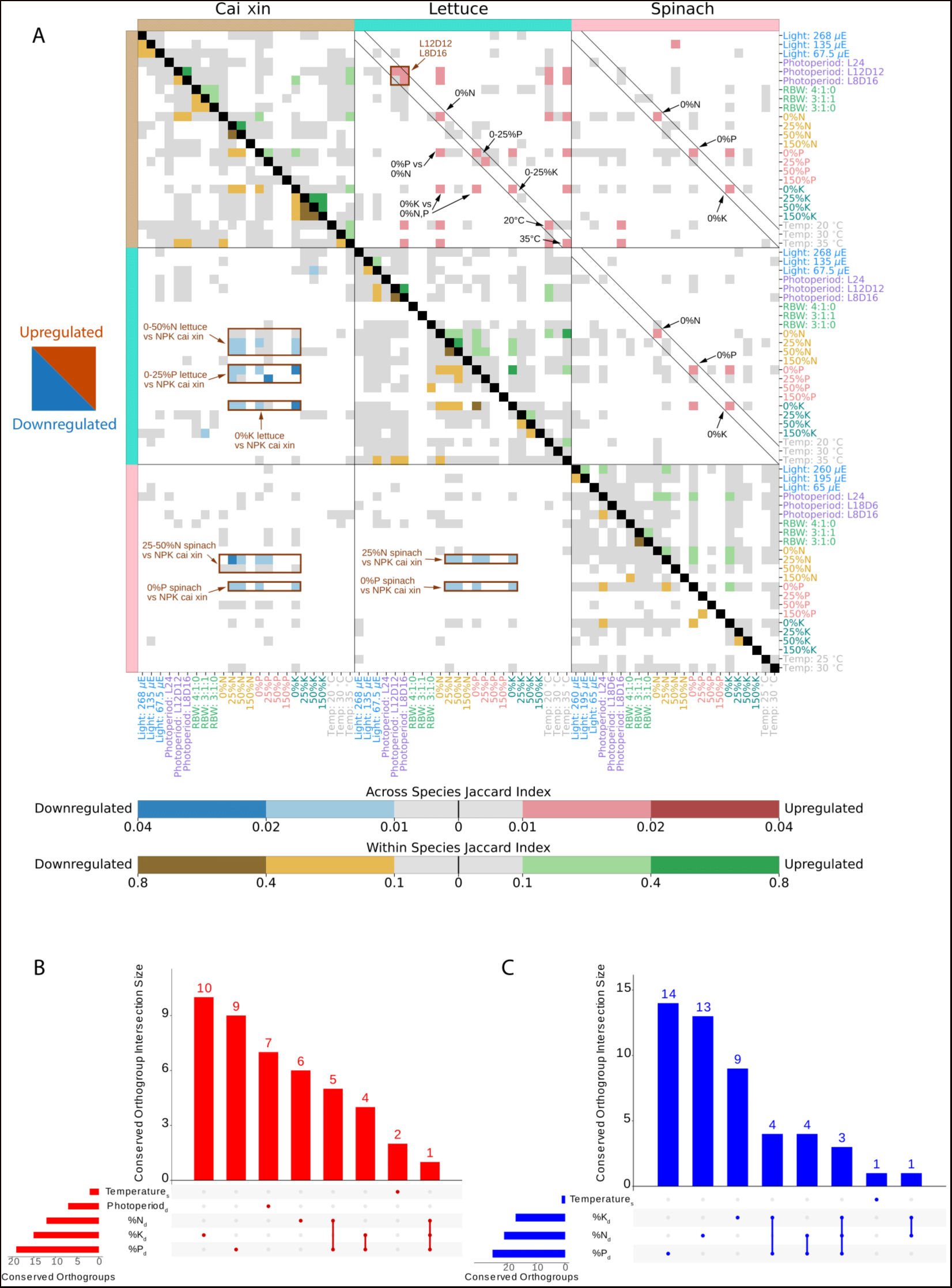
Conservation analysis of transcriptomic stress responses. A) The heatmap shows the conservation of differentially up- (upper, right triangle) and down-regulated (lower, left triangle) genes (and orthogroups) across the three species in the various stress conditions. For the across species analysis, Jaccard index between 0.01 - 0.02 (light red) and 0.02 - 0.04 (dark red) was computed between the upregulated orthogroups of two stress conditions. Similarly, Jaccard indices between 0.01 - 0.02 (light blue) and 0.02 - 0.04 (dark blue) indicate similarities between the downregulated orthogroups of two stress conditions. For within species analysis, Jaccard index between 0.1 - 0.4 (light green) and 0.4 - 0.8 (dark green) was computed between the upregulated DEGs of two stress conditions. Similarly, Jaccard indices between 0.1 - 0.4 (light brown) and 0.4 - 0.8 (dark brown) indicate similarities between the downregulated DEGs of two stress conditions. Gray cells represent Jaccard indices of 0 to 0.01 and 0 to 0.1 for across and within species analyses, respectively. White cells indicate no significance (p-adj > 0.05) for the Jaccard index computed between two stress conditions. Upset plots for B) upregulated and C) downregulated conserved orthogroups across species. The x-axis indicates the different stress condition combinations, while the y-axis indicates the number of orthogroups in a given combination. Subscripts d and s represent deficient and surplus, respectively.

To identify gene families that might be important for stress responses, we identified orthogroups that demonstrated conserved responses across all species under the deficient and/or surplus stress conditions (Figures 4B-C). In complement with Figure 4A’s observations, N, P and K deficiencies stress conditions encompassed the largest numbers of stress condition-specific significantly conserved orthogroups, where 6, 9 and 10 upregulated, and 13, 14 and 9 downregulated orthogroups were identified, respectively. Notably, some conserved orthogroups show up- and/or down-regulation in more than one surplus and/or deficient stress conditions, as seen for N and P deficiencies (5 up- and 4 down-regulated orthogroups), K and P deficiencies (4 up- and 4 down-regulated orthogroups) and N, P, and K deficiencies (1 up- and 3 down-regulated orthogroups) (Figures 4B-C). The genes found in these 89 orthogroups comprise prime candidates to better understand the responses of the species to the different stress conditions (Table S12).

To better understand the functions of the conserved upregulated genes, we performed a literature search of their best BLAST hits from Arabidopsis (Table S12). Various genes have already been reported to be involved in N, P or K deprivation, such as growth regulating factor (Lantzouni et al. 2020) and R2R3 MYB transcription factor (Gaudinier et al. 2018), a calcium-dependent protein kinase (Qin et al. 2020; Liu et al. 2021; Adavi and Sathee 2024) and other signaling components, like the purple acid phosphatase *AtPAP12* (Wang et al. 2014). Glyceraldehyde-3-phosphate dehydrogenase 1 (*GAPCp1*) was reported as an important determinant in the carbon and nitrogen metabolism by connecting glycolysis with other pathways, such as serine biosynthesis, or the ammonium assimilation pathway (Anoman et al. 2015). Interestingly, uridine kinase-like 2 (*UKL2*), an enzyme involved in the uridine salvage pathway (Chen and Thelen 2011), was upregulated during P deficiency. Salvage pathways recover bases and nucleosides that are formed during degradation of RNA and DNA. The salvaged products can then be converted back into nucleotides, which could be particularly important during nutrient starvation (Chen and Thelen 2011). Glutamate dehydrogenase 3 (*GAD3*), which catalyzes a reversible reaction that synthesizes or deaminates glutamate, may play a role in the control of nitrogen metabolism in leaf development (Marchi et al. 2013), while the other members of the GAD family also facilitate acclimation of Arabidopsis to P starvation by GABA shunt upregulation (Benidickson et al. 2023). Apart from the starvation response, in shorter photoperiod, all three species showed upregulation of a gene corresponding *ATNTT2*, which shows necrotic lesions during shrefort photoperiod (Reinhold et al. 2007). Thus, the genes in this list constitute valuable targets for studying how plants respond to abiotic stresses.

### Stress-responsive gene regulatory networks are conserved across species

To understand the conserved relationship between species, conserved TF-target edges were identified between 2 or more species, and permutation analysis was performed to determine the significance of the conserved TF-target edges. From the GRN of cai xin, 1071 TF-target edges were found to be conserved with the GRN of lettuce/spinach, and 19 TF-target edges were conserved with the GRNs of all three species (Figure 5A, Tables S13-S15). The permutation analysis revealed that the TF-target edges were significantly conserved (p-adj < 0.05) between 2 or more species, especially between cai xin and spinach, and lettuce and spinach, where enrichment scores of 1.24 (i.e., 24% more edges than expected by random) and 1.27 were computed, respectively (Figure 5B).

**Figure 5.**
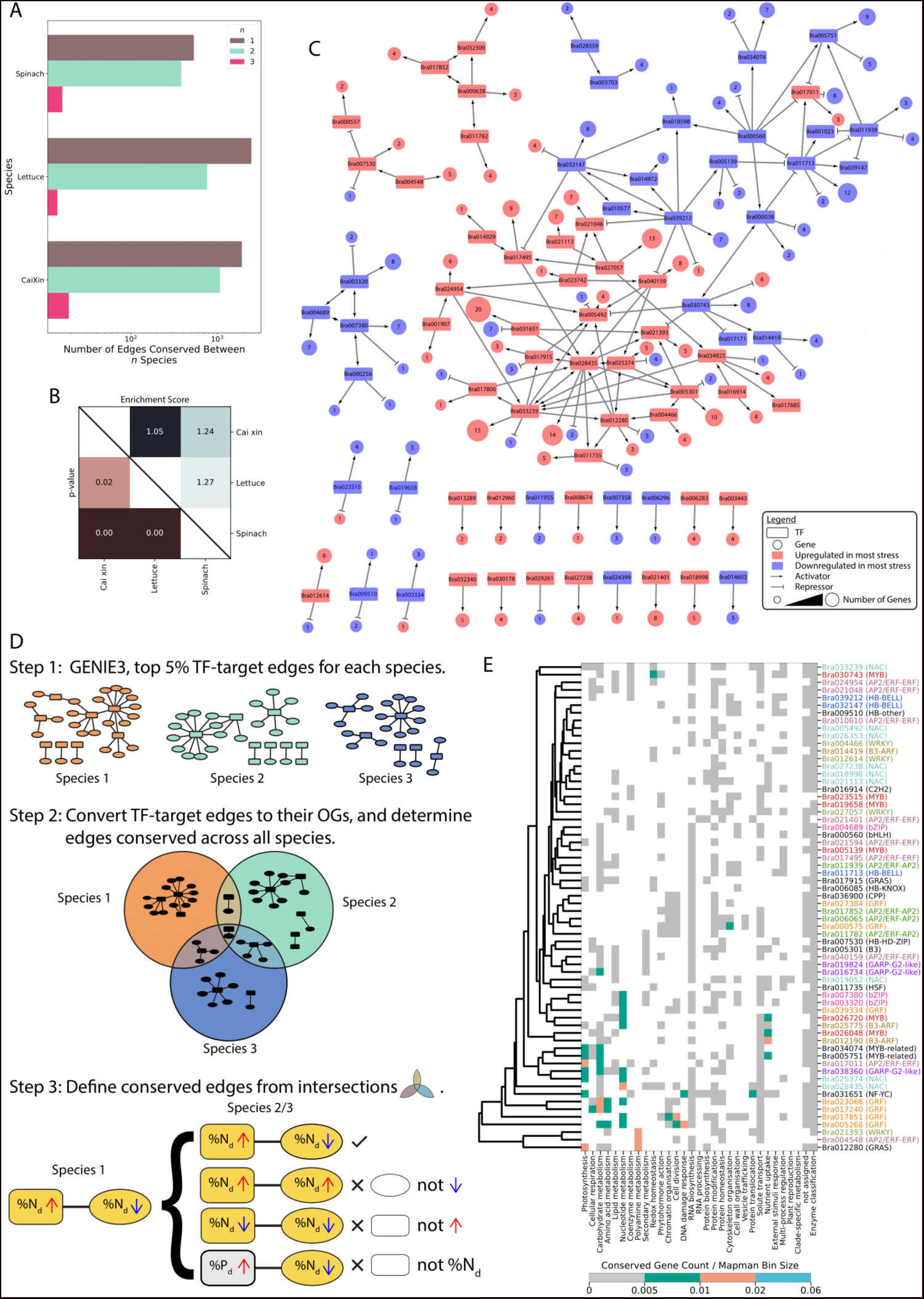
Construction and comparison of gene regulatory networks for cai xin, lettuce, and spinach. A) The number of differentially expressed TF-target edges (leading top 5% by rank of weights) of cai xin, lettuce and spinach, categorized by *n* of 1 (brown bars), 2 (green bars) and 3 (pink bars). Brown bar (*n* = 1) represents edges exclusive to the respective species, green bar (*n* = 2) represents conserved edges between the respective species and one other species, and pink bar (*n* = 3) represents conserved edges between the respective species and all other species. B) The adjusted p-values (lower right triangle) and enrichment scores (upper right triangle) of the similarity of the GRNs across the three species. C) Gene regulatory network for potassium deficiency of cai xin. Rectangular and circular nodes represent TFs and genes (targets), respectively. The size of the circular nodes is indicative of the number of genes a TF regulates, where larger sizes represent a larger group of genes. Delta and T arrows indicate activators and repressors, respectively. Red color represents upregulation in the stress condition-specific GRN, and blue represents downregulation. D) Schematic work flow of the identification of conserved TF-target edges. Step 1, GENIE3 is employed to generate GRN of each species, and the top 5% (by the ranks of non-zero weight) TF-target edges were used for subsequent steps. Step 2, all TF-target edges were mapped to their respective OGs and edges that were conserved across in at least two species were identified. In Step 3, we further refined our analysis by identifying conserved TF-target edges under various conditions. This involved pinpointing edges that were consistently differentially expressed across species and under the same stress condition. E) The heatmap depicts the conserved TFs (columns) and biological functions (rows) they are predicted to regulate. For brevity, only TFs that regulate at least 5 biological functions are shown. The TFs are also color-coded according to their gene symbol for gene symbols that appeared at least 3 times. The cell color indicates the number of targets (and their associated MapMan bins) normalized by the corresponding MapMan bin sizes. White indicates no evidence of TF regulating a given biological function. Gray represents weak regulation. Green, orange, and blue indicate increasing strength of regulation of a TF of a biological function.

Surplus and deficient stress condition-specific GRNs were built with significantly conserved TF-target edges for each species (Tables S13-S15). For the case of K deficiency stress condition GRN of cai xin, some stress condition-specific conserved TFs, such as *Bra031651*, *Bra033239*, and *Bra028435*, were observed to regulate large groups of genes (targets) (Figure 5C). Corresponding networks for different surplus and deficient stress conditions and species were attached in Supplemental Data 4. The work flow of the identification of conserved TF-target edges is depicted in Figure 5D. *Bra031651* is conserved only under K deficiency stress condition, while *Bra033239* and *Bra028435* are also expressed in photoperiod and P deficiencies stress condition, respectively. This observation suggests that certain conserved TFs demonstrate particularly strong regulatory effects under specific stress conditions.

To gain insight into biological functions (MapMan bins) the stress condition-specific conserved TFs are likely to regulate, the distribution of the conserved TFs and their targets’ MapMan bin annotations were investigated (Figure 5E), where only conserved TFs that regulate at least 5 biological functions were depicted for brevity. Results for cai xin revealed that the TFs control multiple biological functions. For instance, *Bra031651*, found to be active under K deficiency stress condition (Figure 5C), regulates multiple biological functions, particularly protein translocation, DNA damage response and photosynthesis. This aligns with *gn_5_74780.1*, an orthologous conserved TFs under K deficiency of lettuce, which also exhibits strong regulation of DNA damage response (Figure S16), and similar distribution across the biological functions. Another instance involves *Bra017011* and *Spov3_chr4.00739* (Figure S17), both of which are active under K deficiency stress condition, and demonstrate strong regulations of photosynthesis and carbohydrate metabolism. Overall, similar distributions across biological functions are observed among the orthologous conserved TFs across all species.

### Limited conservation of regulatory logic of biological pathways

To verify the roles of the conserved transcription factors identified for the three species (Figure 5E and Figures S16-S17), we studied the experimentally verified functions of the *Arabidopsis thaliana* best BLAST hit orthologs (Figure 6A and Table S16). Only nine TFs have conserved responses with their *Arabidopsis thaliana* hits, with the majority of them being found in the phosphate responses. A hit in nitrogen is found for *gn_4_20080.1*, which was both down- and up-regulated under N and P deficiencies stress conditions, respectively, and its best hit, *AT1G13300* has also been shown to respond in nitrogen and phosphate starvation (GO terms). For *Bra016734* and *Bra019824*, which best hits are also *AT1G13300*, show only a match in response to P deficiency. Notably, *Bra006085* and *gn_6_20120.1*, which are differentially expressed under both N and photoperiod deficiencies stress conditions, the expression of their best hits, *AT5G11060*, has only been shown to be regulated by light. The mismatches between the conserved TFs and their *Arabidopsis thaliana* best hits suggested the possibility that these hits might have different functions, or have not yet been studied under certain stresses.

**Figure 6.**
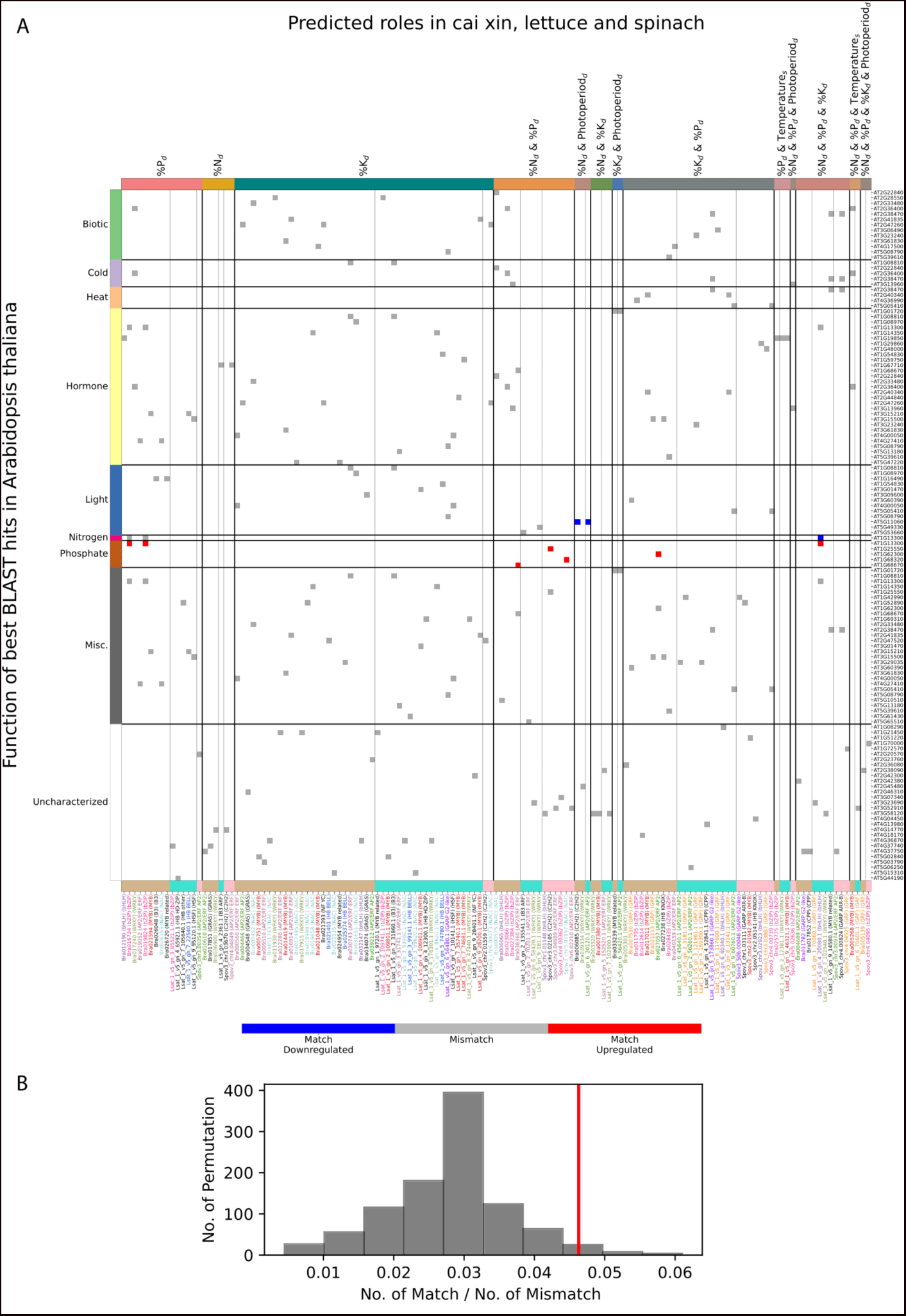
Comparison of stress-responsive transcription factors in lettuce, spinach and cai xin to corresponding experimentally verified genes in *Arabidopis thaliana*. A) The heatmap indicates the conserved transcription factors (columns) of cai xin (tan), lettuce (turquoise) and spinach (pink), and their corresponding *Arabidopsis thaliana* best BLAST hits (rows). The genes are grouped by the stress conditions at which the TFs were differentially expressed, and the different stress conditions and combinations are color-coded (top column colors). The stress conditions include P, N, K, photoperiod deficiencies (subscript d), and surplus (subscript s) temperature. The corresponding best hits are grouped according to their functions (row colors) obtained from research papers. Red and blue cells indicate cases where the reported function of Arabidopsis TF corresponds to the same predicted role in our experiments (matches), while gray squares indicate mismatches. B) The observed ratio of matches to mismatches (red line) to the permuted ratio of matches to mismatches (gray histogram).

To determine whether the function of the TFs are significantly conserved with respect to *Arabidopsis thaliana*, permutation analysis was performed. The results demonstrated conservation to be significant (p-adj = 0.001, Figure 6B), whereby the observed proportion of matches is higher than the permuted matches (enrichment score = 1.67). Therefore, while there is significance, the conservation is mainly due to conserved P and light responses.

## Conclusion and Discussion

Hydroponic cultivation is increasingly favored globally for its efficient resource management and the production of high-quality food. Traditional soil-based agriculture faces numerous obstacles, including urbanization, natural disasters, climate change, and the detrimental impact of excessive chemical and pesticide use, which diminishes soil fertility (Sharma et al. 2019). Our studies show that, although plant growth often does not surpass control conditions, it is possible to reduce the levels of certain nutrients, such as PKN, to 50% of their recommended amounts without significantly impacting growth (Figure 1). Beyond its resource efficiency, hydroponics serves as an invaluable research instrument, enabling the swift experimentation with growth parameters in conditions mimicking the ‘field’. Our research demonstrated this capability by testing 24 growth conditions across three species simultaneously in growth conditions mimicking a hydroponics setup. This approach is critically important, given the low success rate in translating growth-promoting genes from models like *Arabidopsis thaliana* to crops; of 1,671 genes tested in maize, only 22 (1.3%) yielded promising leads for further development (Simmons et al. 2021; Inzé and Nelissen 2022). Unlike field experiments where environmental variables remain uncontrolled, hydroponics offers the flexibility to alter these parameters on the target crop in a parallelized manner, thereby enhancing the throughput, reproducibility and reliability of research findings. Furthermore, since the metabolomic composition of crops can be modulated by the growth conditions (Figure 3) (Hanson et al. 2016), hydroponics could offer an avenue to align the nutrient content of our crops to our needs.

In recent years, nutrient deficiencies have emerged as significant threats to crop growth, production, food safety, and quality (Neset and Cordell 2012; Shahzad et al. 2014). Prior research predominantly explored the mechanisms and signaling pathways that model plants, such as Arabidopsis and rice, employ to maintain homeostasis during individual nutrient shortages (Fan et al. 2021). These studies have enriched our understanding of the genes crucial for mineral nutrient balance under such deficiencies. Through molecular biology, genetics, and omics techniques, key regulators of nitrogen (N), phosphorus (Pi), zinc (Zn), and iron (Fe) absorption and equilibrium in *Arabidopsis thaliana* and rice have been pinpointed during mineral scarcities (Kobayashi and Nishizawa 2012; Park et al. 2014; Bouain et al. 2018; Yang et al. 2018). However, gene expression analyses, despite their value, often identify thousands of differentially expressed genes (Figure 2), posing challenges in distinguishing vital survival genes from those that are merely secondary stress responses. This issue can be mitigated by comparative research, prioritizing genes with consistent expression patterns across species (Julca et al. 2023). For instance, conserved co-expression modules likely denote groups of truly functionally interconnected genes (Movahedi et al. 2011; Mutwil et al. 2011; Hansen et al. 2014). Our findings corroborate this approach, showing a high enrichment of genes essential for phosphate deprivation survival between cai xin and lettuce (Table 1, Figure 2H). Furthermore, we observed a unified response mechanism to NPK depletion across species (Figure 4), hinting at the potential for engineering resilience to these deficiencies by modifying the activity of the shortlisted genes.

Stress adaptation mechanisms are orchestrated at the transcriptional level by transcription factors (TFs), which lead to the accumulation of stress-responsive cellular factors (Ramanjulu and Bartels 2002; Manna et al. 2021), highlighting TFs as pivotal targets for genetic engineering. Leveraging the power of comparative transcriptomics and the parallel nature of our stress experiments, we have devised a novel pipeline that merges traditional regression methods with comparative genomics to construct conserved gene regulatory networks (Figure 5C). Our findings demonstrate significant conservation of these networks across species, delineating the biological pathways influenced by these TFs (Figure 5E), yet reveal a notable but limited conservation of the functions attributed to Arabidopsis orthologs, with exception of phosphate and nitrogen deprivation (Figure 6). This partial conservation may be attributed to the species-specific divergence of gene functions, a phenomenon also noted in animal studies (Berthelot et al. 2018). Additionally, morphological distinctions could influence the impact of certain genes. For instance, the *SAMBA* gene, a negative regulator of cell cycle progression, boosts leaf growth in Arabidopsis through enhanced cell division upon inactivation (Eloy et al. 2012), but in maize, its mutation leads to reduced growth due to possibly excessive cell division during later development stages (Gong et al. 2022a). This indicates that some growth-regulatory networks are exclusive to eudicots and absent in monocots, such as the PEAPOD-KIX-TOPLESS repressor complex (Schneider et al. 2021), which limits growth in various eudicot organs (Naito et al. 2017; Cookson et al. 2022), but is not found in grasses (Schneider et al. 2021). Furthermore, while mutations in *DA1* and *BIG* BROTHER genes result in larger organs in Arabidopsis (Chen et al. 2021), similar mutations in maize do not produce growth-related phenotypes (Gong et al. 2022b), despite gene conservation. However, we cannot exclude the possibility that the roles of Arabidopsis TFs in the stresses examined in this study have not been fully explored, as negative findings often go unreported due to publication biases against non-positive results (Nimpf and Keays 2020). Fortunately, there is a growing acknowledgment of the value of reporting such “lost in translation” findings, as evidenced by recent publications (Gong et al. 2022b). The extensive availability of public data across diverse stresses and species presents a unique opportunity to accurately identify genes that confer stress resilience (Julca et al. 2023), emphasizing the importance of cross-species analyses and the potential for translational insights into stress tolerance mechanisms.

## Acknowledgments

We are grateful to Singapore Food Agency for sponsoring this research (grant SFS_RND_SUFP_001_05). We would like to thank ComCrop for providing us with the lettuce and cai xin seeds, and to ArtisanGreen for providing us with spinach seeds. Finally, we would like to acknowledge Prof. Oliver Mueller-Cajar for his help with the experimental design. We would like to thank Zahin Mohd Ali and Qiao Wen Tan for their help with setting up the stress.plant.tools database.

## Author contributions

J.M.L performed the computational analyses, J.C.G and D.M.A were involved in the experimental design and execution, M.M supervised the project.

## Supplementary Figures

Figure S1. Cluster Map for cai Xin.

Figure S2. Cluster Map for lettuce.

Figure S3. Cluster Map for spinach.

Figure S4. Pathway enrichment analysis (second-level) for Spinach. Hierarchical clustering of the biological pathways and stress conditions is computed according to Jaccard distance.

Figure S5. Pathway enrichment analysis (second-level) for lettuce.

Figure S6. Pathway enrichment analysis (second-level) for cai xin.

Figure S7. Nicotinamide concentrations and gene expression levels (TPM) of genes involved in nicotine biosynthesis pathway.

Figure S8. Nicotinic acid concentrations and gene expression levels (TPM) of genes involved in nicotine biosynthesis pathway.

Figure S9. Pantothenic acid concentrations and gene expression levels (TPM) of genes involved in phosphopantothenate biosynthesis I pathway.

Figure S10. Pyridoxal concentrations and gene expression levels (TPM) of genes involved in pyridoxal 5’-phosphate biosynthesis II pathway.

Figure S11. Pyridoxine concentrations and gene expression levels (TPM) of genes involved in pyridoxal 5’-phosphate biosynthesis II pathway.

Figure S12. Riboflavin concentrations and gene expression levels (TPM) of genes involved in flavin biosynthesis I (bacteria and plants) pathway.

Figure S13. Thiamine concentrations and gene expression levels (TPM) of genes involved in thiamine diphosphate biosynthesis IV (eukaryotes) pathway.

Figure S14. Conserved lettuce TFs that are controlling >4 mapman bins. Figure S15. Conserved spinach TFs that are controlling >4 mapman bins.

## Supplementary Tables

Table S1. Fresh weight data for all plants

Table S2. Description of the RNAseq experiments (number of reads, % reads mapped, experimental annotation)

Table S3. Deseq2 table for for lettuce

Table S4. Deseq2 table for for cai xin

Table S5. Deseq2 table for for spinach

Table S6. Vitamin levels

Table S7. Correlation between vitamin levels and gene expression

Table S8. Conservation of stress responses for cai xin

Table S9. Conservation of stress responses for lettuce

Table S10. Conservation of stress responses for spinach

Table S11. Conservation of stress responses for spinach

Table S12. IDs of genes found in the conserved Ogs

Table S13. Conserved GRN for caixin

Table S14. Conserved GRN for lettuce

Table S15. Conserved GRN for spinach

Table S16. Experimentally verified functions of Arabidopsis genes and their best blast hits from the three species

## Supplementary Data

Supplemental Data 1. Expression matrices of cai xin

Supplemental Data 2. Expression matrices of lettuce

Supplemental Data 3. Expression matrices of spinach

Supplemental Data 4. Gene regulatory networks for cai xin, lettuce and spinach

