## Supplementary material for "Parallel comparison reveals conservation and re-wiring of abiotic stress response networks across three plant species": Figure S1

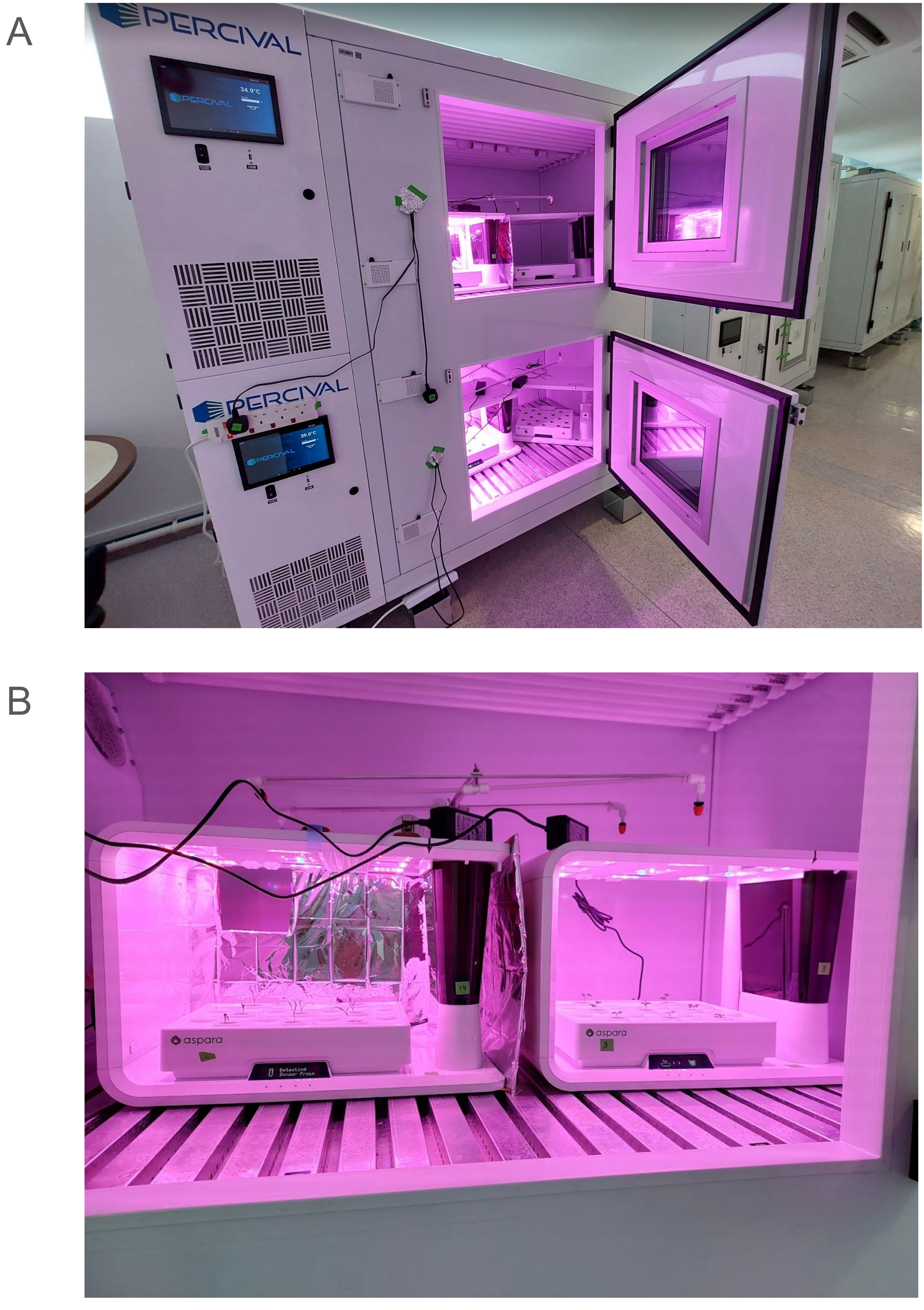


Figure S1. Experimental setup. A) The environmental stresses were performed in the Percival chamber model PGC-9 series (Percival Scientific, Inc, Perry, US). B) All plants were grown in Aspara hydroponic unit Aspara® Nature+ Smart Growers (Growgreen Ltd. Hong Kong).
