## Supplementary material for "Parallel comparison reveals conservation and re-wiring of abiotic stress response networks across three plant species": Figure S2

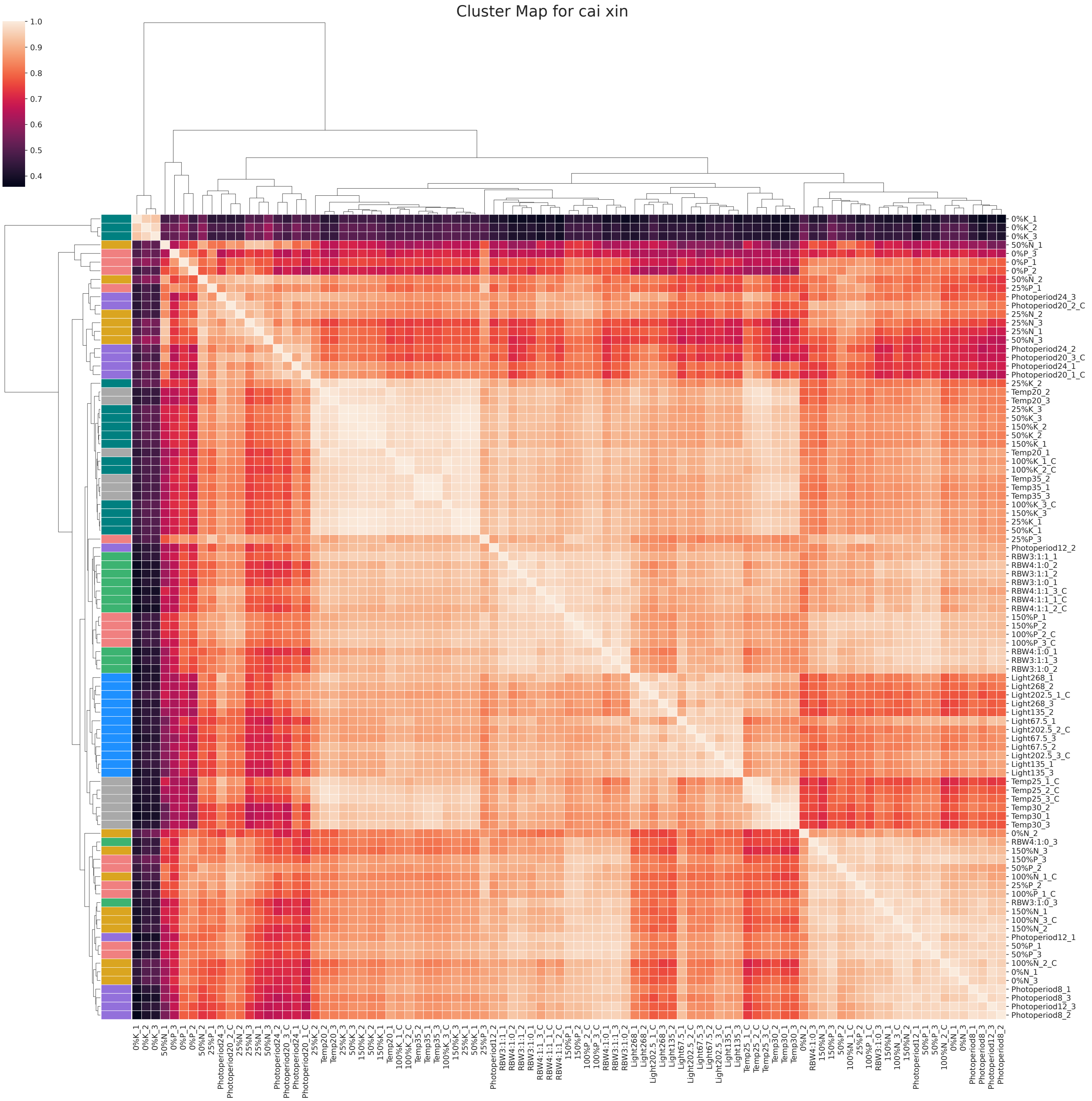


Figure S2. Pearson correlation coefficient between all the samples and stress conditions of the cai xin stress experiments based on the TPM gene expression values. Hierarchical clustering of the all the samples indicates similarity between the samples and stress conditions.
