## Supplementary material for "Parallel comparison reveals conservation and re-wiring of abiotic stress response networks across three plant species": Figure S3

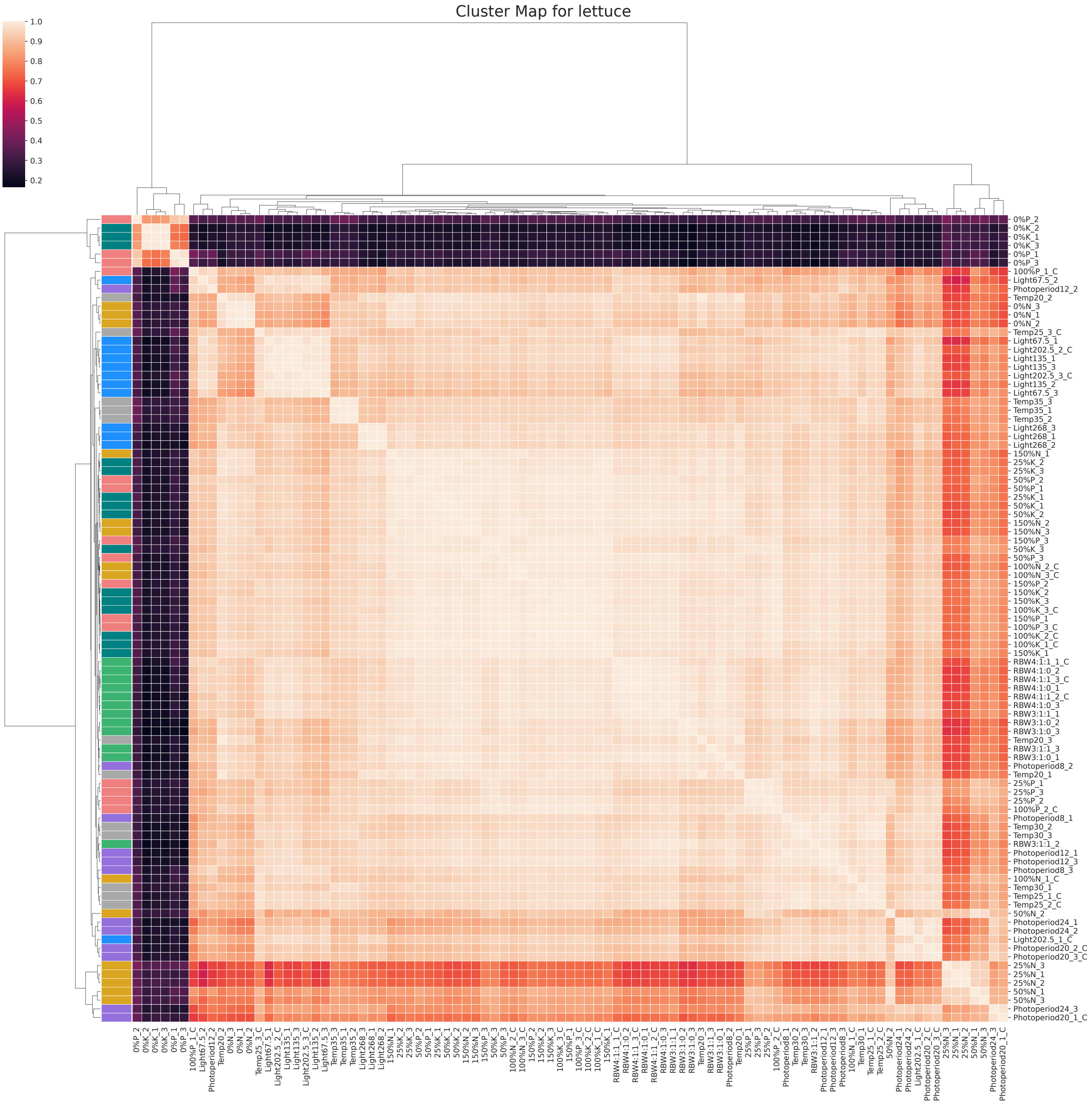


Figure S3. Pearson correlation coefficient between all the samples and stress conditions of the lettuce stress experiments based on the TPM gene expression values. Hierarchical clustering of the all the samples indicates similarity between the samples and stress conditions.
