## Supplementary material for "Parallel comparison reveals conservation and re-wiring of abiotic stress response networks across three plant species": Figure S5

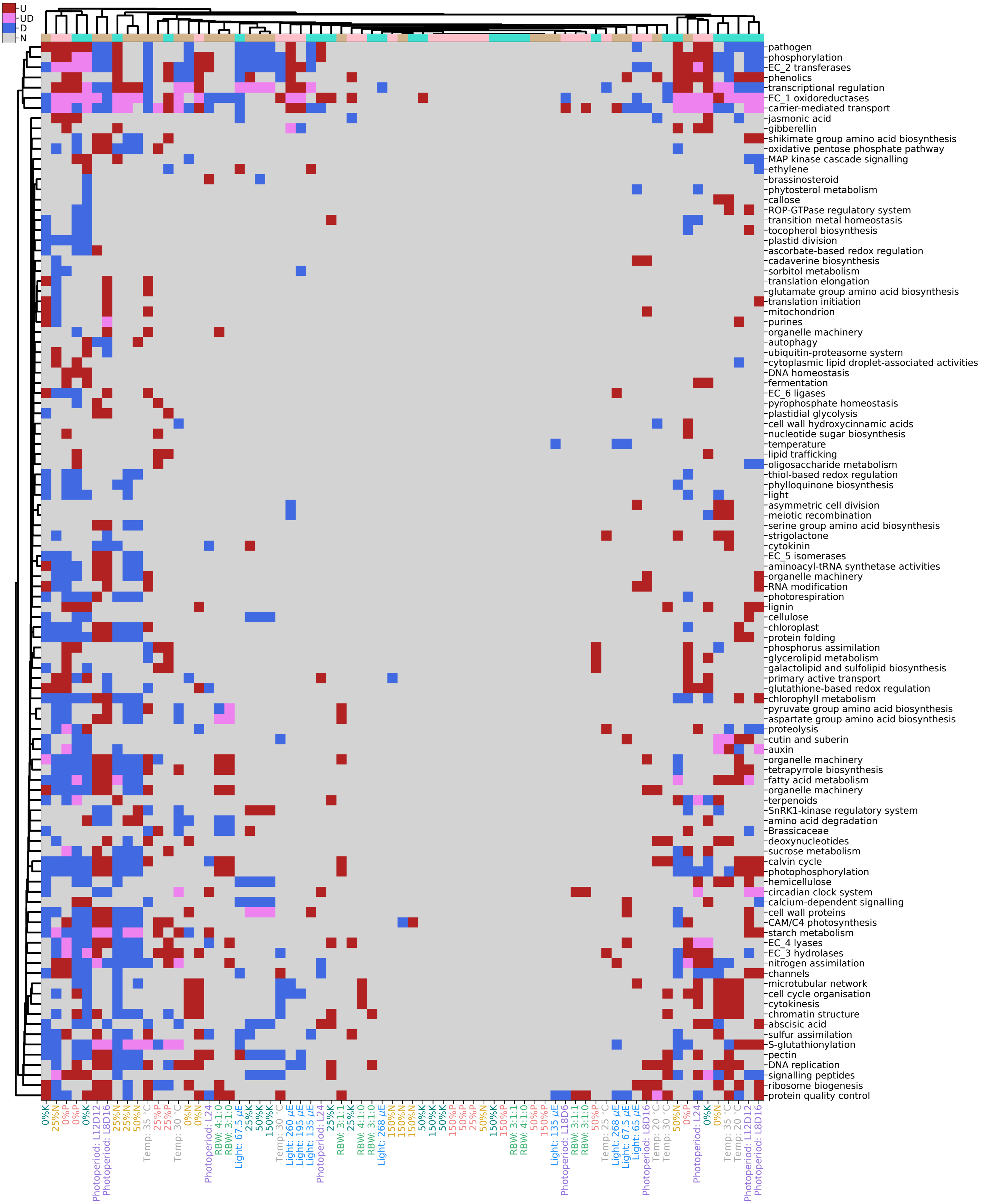


Figure S5. Significantly (p-adj < 0.05) up- (red), down-regulated (blue) or both (pink) level 2 biological functions (Mapman sub-bins) for the different stress conditions and species. The different stress conditions are in columns, while the biological pathways are in rows. The labels of the stress condition are colored according to their stress experiments. Similarities between the responses were highlighted by clustering the columns and biological functions across all three species, and the columns are color-coded such that tan, turquoise and pink represent cai xin, lettuce and spinach, respectively.
