## Supplementary material for "Parallel comparison reveals conservation and re-wiring of abiotic stress response networks across three plant species": Figure S6

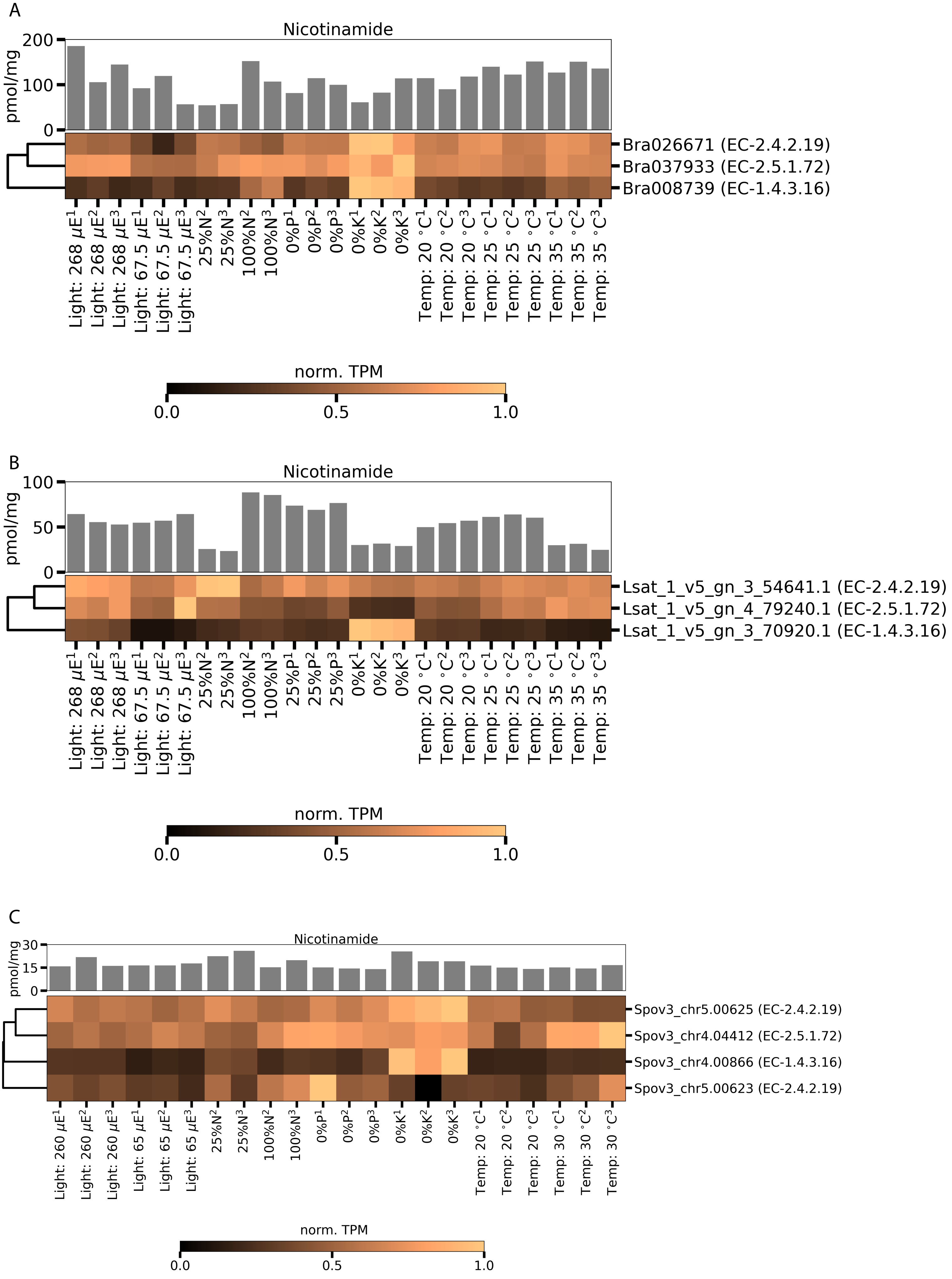


Figure S6. Nicotinamide levels (in pmol/mg) and row-normalized TPM of the first four steps biosynthetic enzymes for various stress conditions of lettuce are represented by a bar chart and heatmap, respectively.
