## Supplementary material for "Parallel comparison reveals conservation and re-wiring of abiotic stress response networks across three plant species": Figure S8

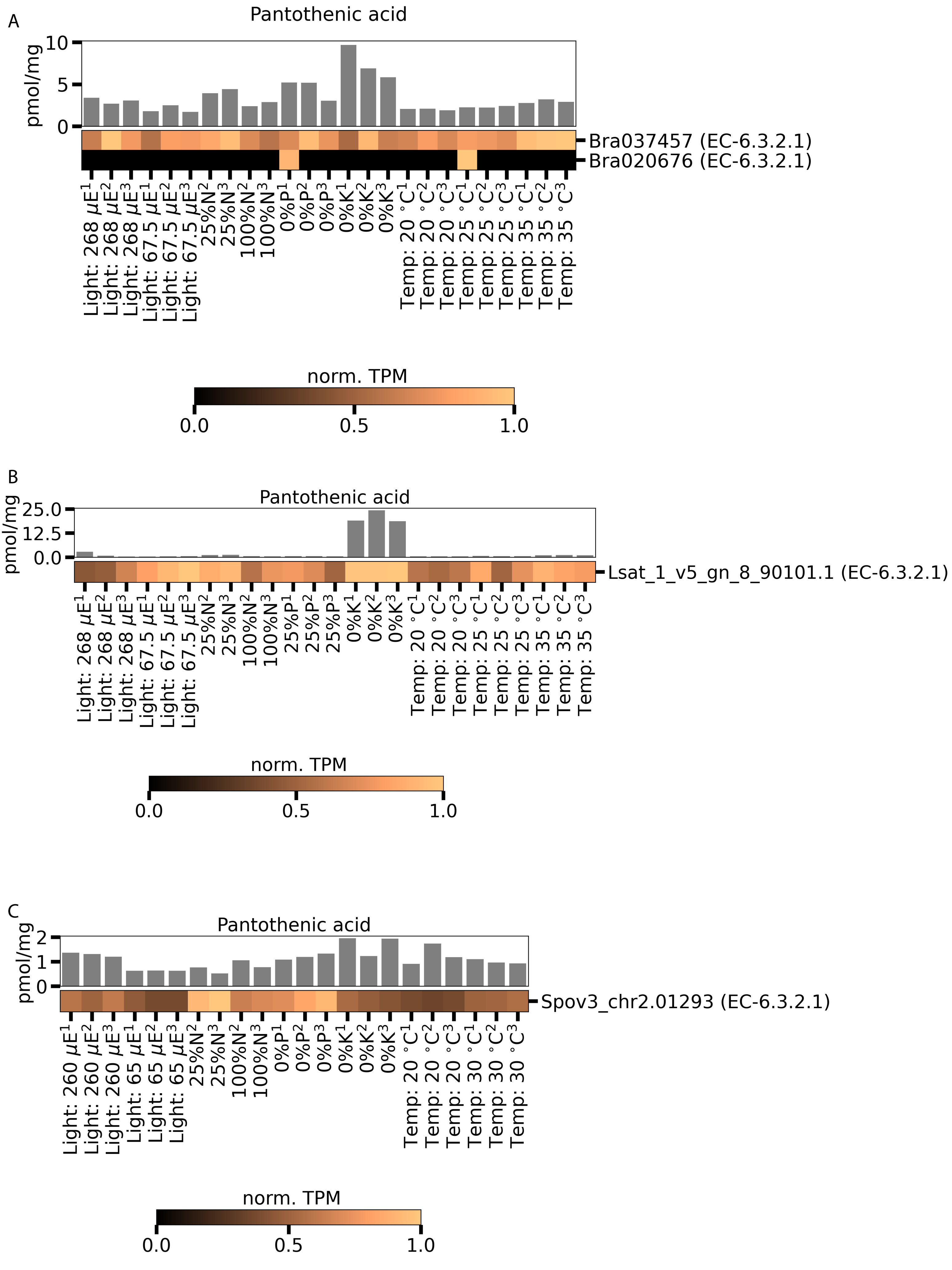


Figure S8. Pantothenic acid levels (in pmol/mg) and row-normalized TPM of the last second step biosynthetic enzymes for various stress conditions of lettuce are represented by a bar chart and heatmap, respectively.
