## Supplementary material for "Parallel comparison reveals conservation and re-wiring of abiotic stress response networks across three plant species": Figure S11

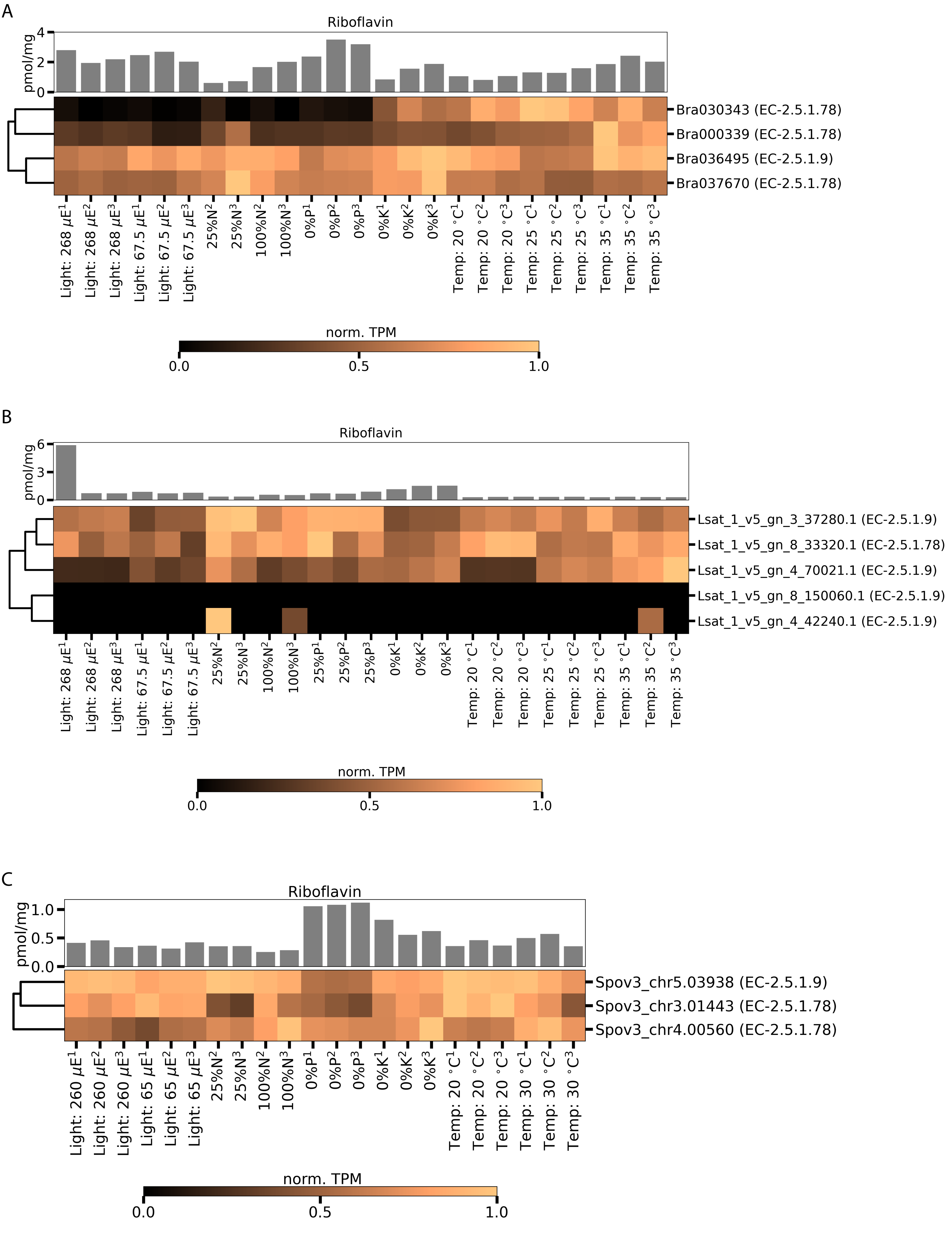


Figure S11. Riboflavin levels (in pmol/mg) and row-normalized TPM of the last third and fourth steps biosynthetic enzymes for various stress conditions of A) cai xin, B) lettuce, and C) spinach are represented by bar chart and heatmap, respectively.
