## Supplementary material for "Parallel comparison reveals conservation and re-wiring of abiotic stress response networks across three plant species": Figure S12

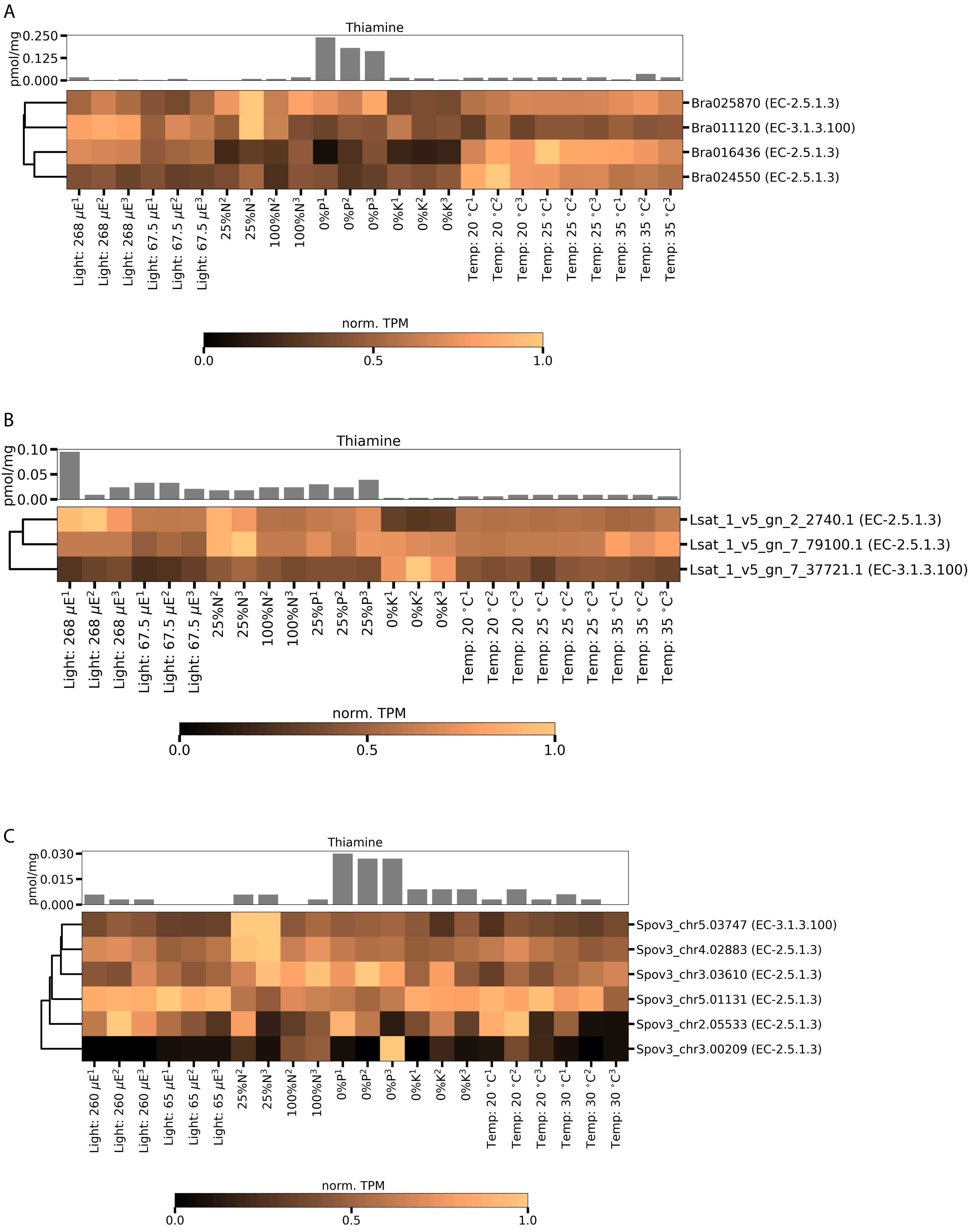


Figure S12. Thiamine levels (in pmol/mg) and row-normalized TPM of the first two steps biosynthetic enzymes for various stress conditions of A) cai xin, B) lettuce, and C) spinach are represented by bar chart and heatmap, respectively.
