## Supplementary material for "Parallel comparison reveals conservation and re-wiring of abiotic stress response networks across three plant species": Figure S13

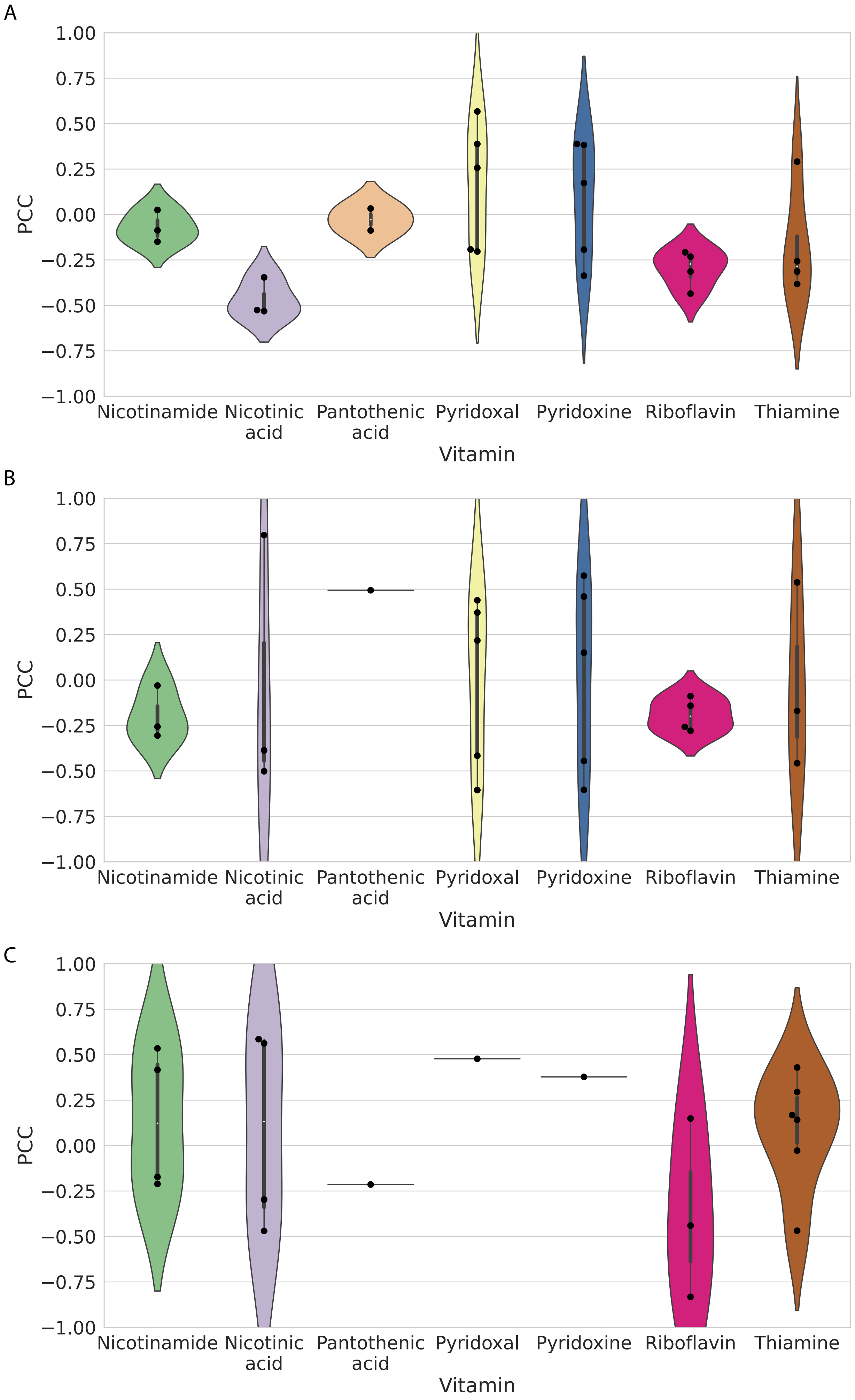


Figure S13. Violin plots depict Pearson correlation coefficient (PCC), capturing the similarity of expression levels of enzymes and the corresponding vitamins for (A) cai xin, (B) lettuce, and (C) spinach.
