## Supplementary material for "Parallel comparison reveals conservation and re-wiring of abiotic stress response networks across three plant species": Figure S15

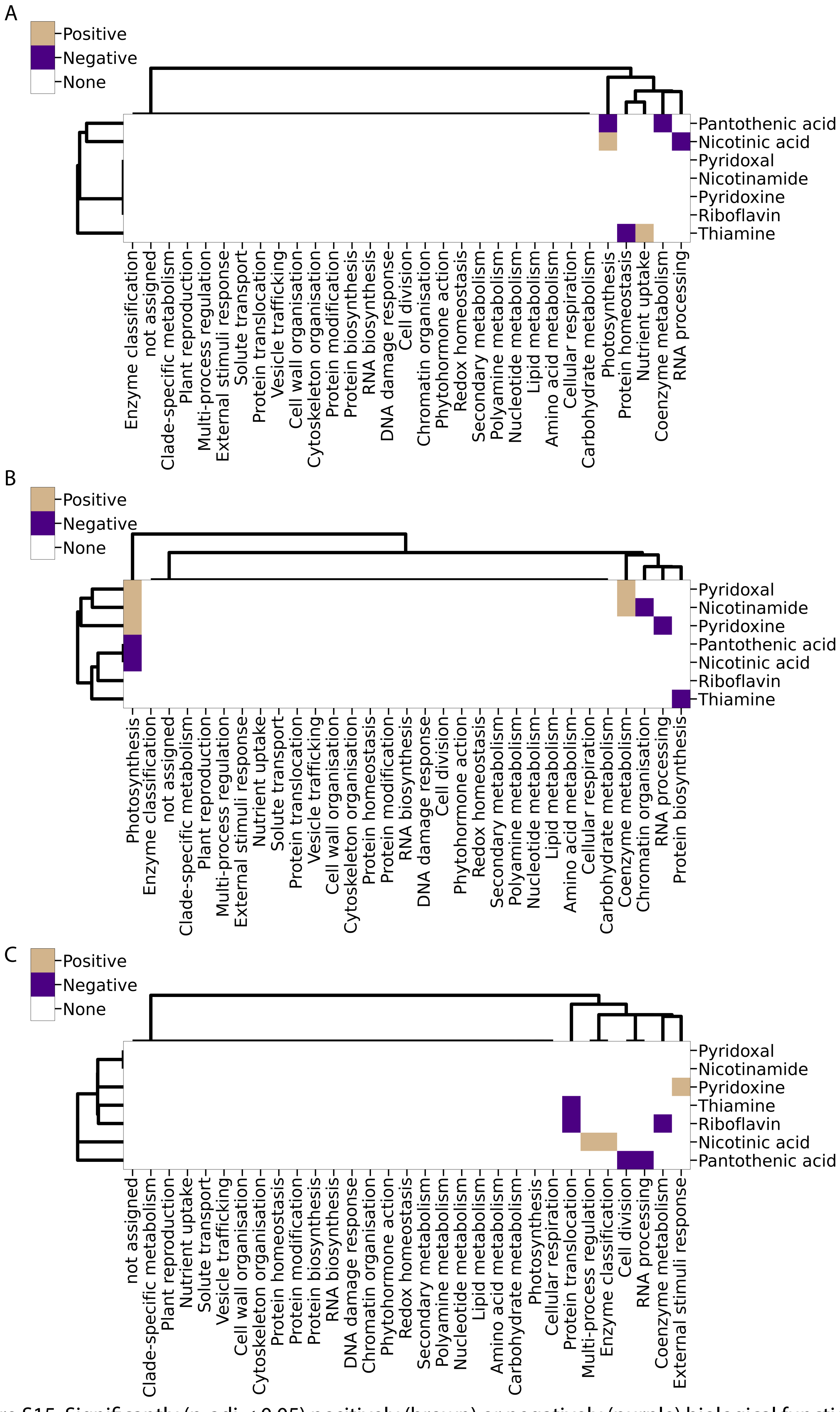


Figure S15. Significantly (p-adj < 0.05) positively (brown) or negatively (purple) biological functions for the different vitamins of (A) cai xin, (B) lettuce and (C) spinach based on the leading 50 positively and negatively PCC correlated genes. White color indicates non-significant correlations between the biological functions and the vitamins. The different biological functions are in columns, while the vitamins are in rows. Similarities between the responses were highlighted by clustering the columns and rows.
