## Supplementary material for "Parallel comparison reveals conservation and re-wiring of abiotic stress response networks across three plant species": Figure S16

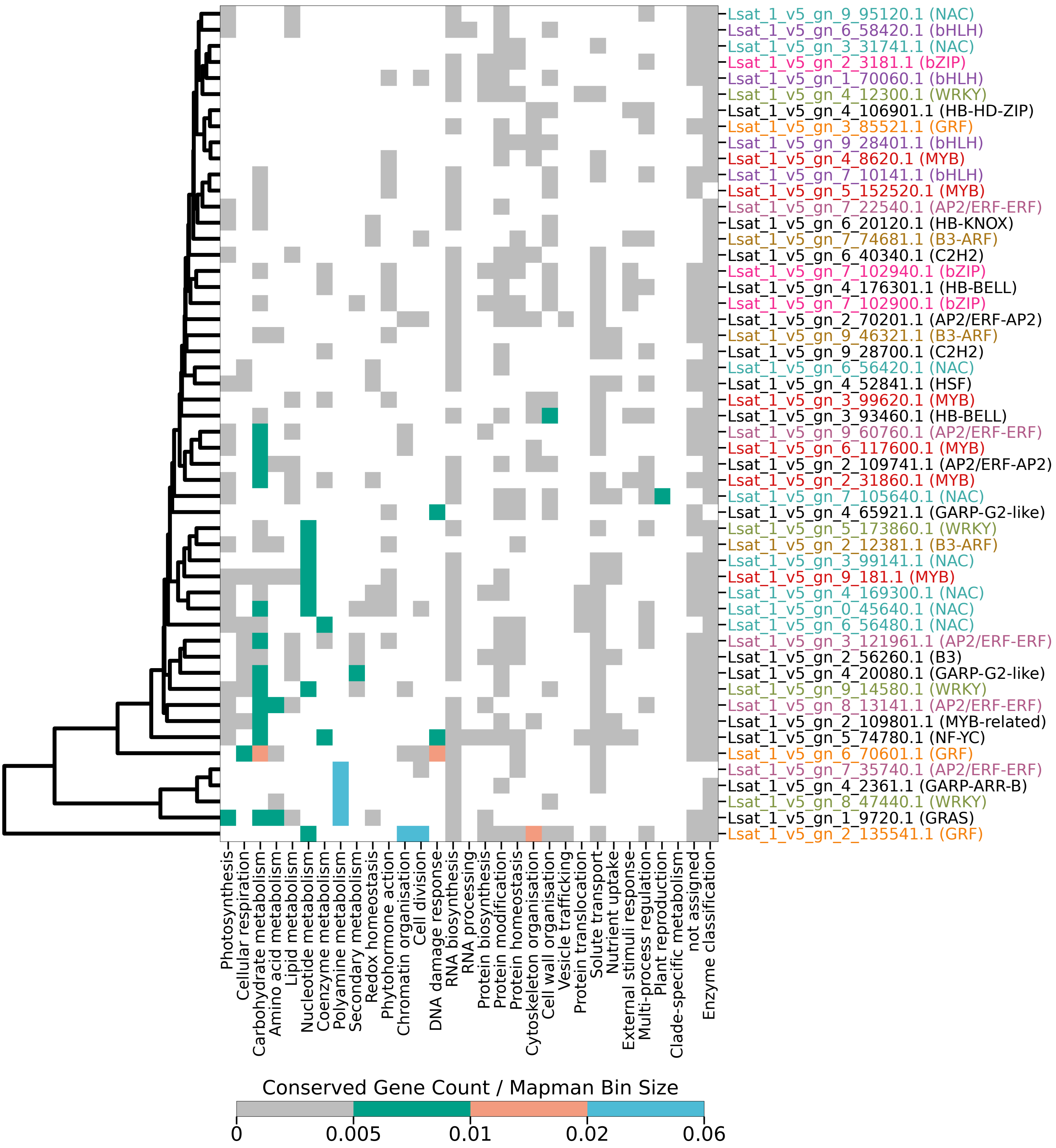


Figure S16. The heatmap depicts the conserved TFs (columns) and biological functions (rows) they are predicted to regulate. For brevity, only TFs that regulate at least 5 biological functions are shown. The TFs are also color-coded according to their gene symbol for gene symbols that appeared at least 3 times. The cell color indicates the number of targets (and their associated MapMan bins) normalized by the corresponding MapMan bin sizes. White indicates no evidence of TF regulating a given biological function. Gray represents weak regulation. Green, orange, and blue indicate increasing strength of regulation of a TF of a biological function.
